# Computational inference of eIF4F complex function and structure in human cancers

**DOI:** 10.1101/2023.08.10.552450

**Authors:** Su Wu, Gerhard Wagner

## Abstract

The canonical eukaryotic initiation factor 4F (eIF4F) complex, composed of eIF4G1, eIF4A1, and the cap-binding protein eIF4E, plays a crucial role in cap-dependent translation initiation in eukaryotic cells (1). However, cap-independent initiation can occur through internal ribosomal entry sites (IRESs), involving only eIF4G1 and eIF4A1 present, which is considered to be a complementary process to cap-dependent initiation in tumors under stress conditions (2). The selection and molecular mechanism of specific translation initiation in human cancers remains poorly understood. Thus, we analyzed gene copy number variations (CNVs) in TCGA tumor samples and found frequent amplification of genes involved in translation initiation. Copy number gains in *EIF4G1* and *EIF3E* frequently co-occur across human cancers. Additionally, *EIF4G1* expression strongly correlates with genes from cancer cell survival pathways including cell cycle and lipogenesis, in tumors with *EIF4G1* amplification or duplication. Furthermore, we revealed that eIF4G1 and eIF4A1 protein levels strongly co-regulate with ribosomal subunits, eIF2, and eIF3 complexes, while eIF4E co-regulates with 4E-BP1, ubiquitination, and ESCRT proteins. Using Alphafold predictions, we modeled the eIF4F structure with and without eIF4G1-eIF4E binding. The modeling for cap-dependent initiation suggests that eIF4G1 interacts with eIF4E through its N-terminal eIF4E-binding domain, bringing eIF4E near the eIF4A1 mRNA binding cavity and closing the cavity with both eIF4G1 HEAT-2 domain and eIF4E. In the cap-independent mechanism, α-helix_5_ of eIF4G1 HEAT-2 domain instead directly interacts with the eIF4A1 N-terminal domain to close the mRNA binding cavity without eIF4E involvement, resulting in a stronger interaction between eIF4G1 and eIF4A1.

**Significance Statement:** Translation initiation is primarily governed by eIF4F, employing a “cap-dependent” mechanism, but eIF4F dysregulation may lead to a “cap-independent” mechanism in stressed cancer cells. We found frequent amplification of translation initiation genes, and co-occurring copy number gains of *EIF4G1* and *EIF3E* genes in human cancers. *EIF4G1* amplification or duplication may be positively selected for its beneficial impact on the overexpression of cancer survival genes. The co-regulation of eIF4G1 and eIF4A1, distinctly from eIF4E, reveals eIF4F dysregulation favoring cap-independent initiation. Alphafold predicts changes in the eIF4F complex assembly to accommodate both initiation mechanisms. These findings have significant implications for evaluating cancer cell vulnerability to eIF4F inhibition and developing treatments that target cancer cells with dependency on the translation initiation mechanism.

## Introduction

Translation initiation is the most crucial regulatory step in protein synthesis, where the eukaryotic initiation factor 4F (eIF4F) complex binds to activated mRNA and recruits ribosomes to translate it into a functional protein (1). Cancer cells rely heavily upon eIF4F to drive aberrant protein synthesis that supports their survival, proliferation, and metastasis (3-5). The canonical eIF4F complex contains three core factors: the scaffold protein eIF4G1, the cap-binding protein eIF4E, and the RNA helicase eIF4A1. eIF4G1 has multiple binding domains, which variously can interact with the 5’UTR of mRNA, eIF4E, eIF4A1, the eIF3 complex, PABP1, and Mnk1/2 (6). When eIF4G1 binds to mRNA, its interaction with eIF4E can force proximity between eIF4E and the 7-methylguanosine cap at the 5’ end of mRNAs, stabilizing the cap and eIF4E interaction (7, 8). eIF4E in turn modulates eIF4G1’s ability to stimulate eIF4A1, which unwinds the mRNA secondary structure for the ribosome attachment (9). The eIF3e subunit of the eIF3 complex directly interacts with eIF4G1, as a connection between eIF4G1 and the 40S ribosomal subunit (10, 11).

Overexpression or activation of eIF4F subunits by oncogenic signaling pathways can increase protein synthesis, promoting tumor growth (12). Cancer development involves genetic alterations enabling cell transformation, survival, drug resistance, and metastasis (13, 14). Selective genetic alterations in translation initiation genes are likely critical for tumor adaptation. However, quantification of positive selection in translation initiation across human cancers is lacking, impeding efforts to target eIF4F inhibition effectively.

eIF4F can initiate translation through a cap-dependent mechanism (with eIF4E), or a cap-independent mechanism (without eIF4E) (1, 15). eIF4E in its own right plays a dual role, in translation initiation, and the export of cell cycle gene mRNAs from the nucleus to cytoplasm through the nuclear pore complex (NPC) (16). Nutrient deprivation can cause eIF4E to accumulate in the nucleus in favor of mRNA transport, reducing its availability for eIF4F (17). Imbalanced expression of *EIF4G1* and *EIF4E* genes have been suggested to dysregulate cap-dependent initiation and favor cap-independent initiation in human cancers (18). Factors such as eIF4E binding protein 1 (4E-BP1) can facilitate cap-independent mechanisms under hypoxia (19) by preventing eIF4E from participating in eIF4F. On a molecular level, translation initiation typically requires eIF4A1 to interact with eIF4G1’s HEAT-1 domain. For cap-independent initiation, eIF4A1 must furthermore interact with eIF4G1’s HEAT-2 domain (20, 21). However, cellular pathways triggering cap-independent initiation remain incompletely understood, and structural insight into eIF4F complex assembly for both mechanisms is limited. Addressing these questions is crucial for the development of cancer treatment drugs that target translation initiation.

To understand the importance of eIF4F components, we employed computational methods on large public datasets to investigate the impact of positive selection on eIF4F dysregulation in cancer. By analyzing copy number variation (CNV) and RNA-Seq data from over 10,000 tumors in The Cancer Genome Atlas (TCGA), we found that translation initiation genes often exhibit co-occurring amplification in tumors. We observed a strong correlation between the expression of *EIF4G1* and genes associated with cancer survival, in tumors with *EIF4G1* amplification or duplication (gain). Our analysis of proteomics data from the Cancer Cell Line Encyclopedia (CCLE) and CRISPR loss-of-function screen data from the Cancer Dependency Map (DepMap) revealed that eIF4E and eIF4G1 are essential for cancer cell survival, but co-regulate with different protein complexes. Using Alphafold, we predicted eIF4F structure with and without eIF4G1-eIF4E binding, and observed varying conformations of the HEAT-2 domain. The predicted contacts between HEAT-2 and the eIF4A1 N-terminal domain are consistent with the experimental structure of an eIF4A1/eIF4G complex (22).

## Results

### Co-occurring copy number gain of translation initiation genes in TCGA tumor samples

Although the *EIF4G1* gene is frequently amplified in human cancers and is considered to be a driver gene (18), its frequency of amplification has not been assessed in the context of copy number variations (CNVs) across all genes in human cancers. To address this, we identified 703 genes amplified in at least 5% of tumors (Fig. 1*A* and Table S1). Gene amplification was less frequent than duplication (Fig. S1*A*), as it often occurs through multiple steps following duplication (23). We focused on amplifications over duplications, because amplifications represent more persistent genomic changes under selection pressure (24).

**Figure 1.**
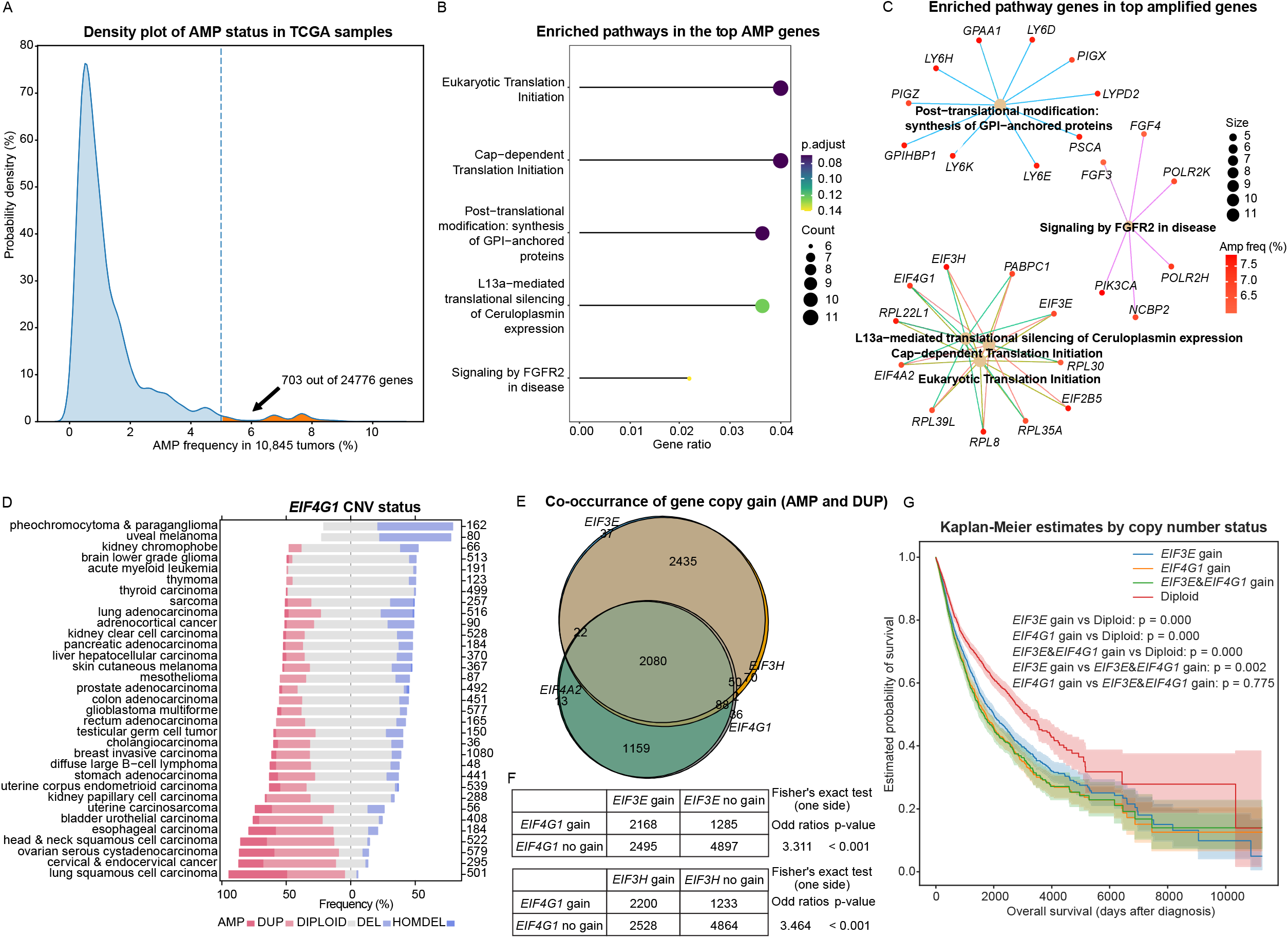
Translation initiation genes frequently co-amplify in TCGA tumor samples. (*A*) The Kernel density estimate plot shows the frequency of cellular gene amplification in tumors across all cancer types in the TCGA database. The highlighted region includes genes amplified in more than 5% of tumors (right of dashed line). (*B*) The dot plot displays pathway enrichment analysis results for the top 703 frequently amplified genes identified in (*A*). Gene ratio represents the ratio of pathway-associated genes to total genes of interest. Dot size indicates the number of genes in each pathway. “p-adjust” is the adjusted p-value using a hypergeometric distribution test for multiple comparisons. (*C*) The star network plot illustrates the genes in the enriched pathways from (*B*). Central node size represents the number of genes in each pathway. Peripheral node color indicates the amplification frequency of corresponding genes. (*D*) The stacked bar plot displays *EIF4G1* copy number variation (CNV) status in different cancer types. (*E*) The Venn diagrams show the co-occurrence of copy number gain (amplification or duplication) for indicated genes in TCGA tumors. (*F*) presents the co-occurrence analysis of copy number gain events for two genes using a contingency table. A positive odds ratio indicates a trend toward co-occurrence. A one-sided Fisher’s exact test was performed to test the null hypothesis of no relationship between the copy number gain events for the two genes. A low p-value suggests that the observed pattern is unlikely due to chance and more likely driven by a specific biological mechanism. (*G*) The KM plot displays the survival probabilities of TCGA patients with cancer based on gene copy gain of *EIF4G1* and/or *EIF3E* (indicated inside each box). The statistical significance of differences was determined by p values from log-rank tests. The shaded areas represent a 95% confidence region for the curve.

Among the 703 most frequently amplified genes in human cancers, we found a significant enrichment of biological functions related to translation initiation, glycosylphosphatidylinositol-anchored protein synthesis, and FGFR2 signaling pathways (Fig. 1*B* and *C*). Initiation factors such as *EIF4G1, EIF3E, EIF3H, EIF4A2, EIF2B5*, and *PABPC1*, and several genes encoding cellular and mitochondrial ribosomal large subunits, were among the most frequently amplified in TCGA cancers (Fig. S1*B*). In addition, we analyzed the correlation of their copy number values across tumor samples. *EIF4G1, EIF4A2*, and *EIF2B5* display strong statistical associations (Fig. S1*C*), likely due to their chromosomal proximity at 3q27. Similarly, we found strong statistical associations between *EIF3E, EIF3H*, and *PABPC1*, likely due to their chromosomal proximity at 8q23.

*EIF4G1, EIF4A2, EIF3E*, and *EIF3H* are frequently duplicated or amplified in most TCGA cancer types, particularly in lung squamous cell carcinoma and head & neck squamous cell carcinoma (Fig 1*D*, Fig. S1*D*, Fig. S2 *A* and *B*). Furthermore, we observed a significant portion of the tumors had copy number gain in both *EIF4G1* and *EIF3E* (Fig 1*E* and Fig. S2*C*), even though they are located on different chromosomes. Fisher’s exact test showed that the co-occurrence of copy number gains in these genes was significantly higher than what would be expected by chance (Fig 1*F* and Fig. S2*D*). Similar co-occurrence of copy number gains was observed between *EIF4G1* and *EIF3H*.

Kaplan-Meier analysis revealed that patients with copy number gains in either *EIF3E* or *EIF4G1* had significantly worse survival probabilities than those with diploid status for both genes in all TCGA cancer types (Fig 1*G* and Fig. S2*E*). Patients with copy number gain in both *EIF3E* and *EIF4G1* had even worse survival probabilities than those with gain in only *EIF3E*. These findings indicate that genes from the eIF4F and eIF3 complexes often exhibit co-occurring copy number gain, which is associated with cancer progression. Moreover, the benefits tumor cells derive from copy number gain in *EIF4G1* may depend on the co-occurring gain of other translation initiation genes.

### A strong correlation between *EIF4G1* expression and cancer survival genes in tumors with *EIF4G1* copy number gain

To determine whether the observed gene amplification is due to positive selection for the tumor-promoting effects of translation initiation genes or simply due to the susceptibility of these loci to amplification (25), we studied the cellular impact of initiation gene amplification on the cellular transcriptome. We identified genes that differentially correlate with *EIF4G1* mRNA expression in TCGA tumors categorized based on their *EIF4G1* CNV statuses and in GTEx healthy tissues.

We divided the correlating genes into five clusters (Fig. 2*A*) and performed pathway enrichment analysis (Fig. 2*B*). We found that the genes in “cluster 4” had strong positive correlations with *EIF4G1* expression in tumors that contained *EIF4G1* copy number gain. However, the strength of these correlations decreased in tumors with *EIF4G1* diploid or deletion, as well as in healthy tissues. In contrast, when we performed similar analyses on *EIF3E* and *EIF3H*, we did not identify any gene clusters that had stronger correlations in tumors with a gain of *EIF3E* or *EIF3H* (Fig. S3*A* to *D*).

**Figure 2.**
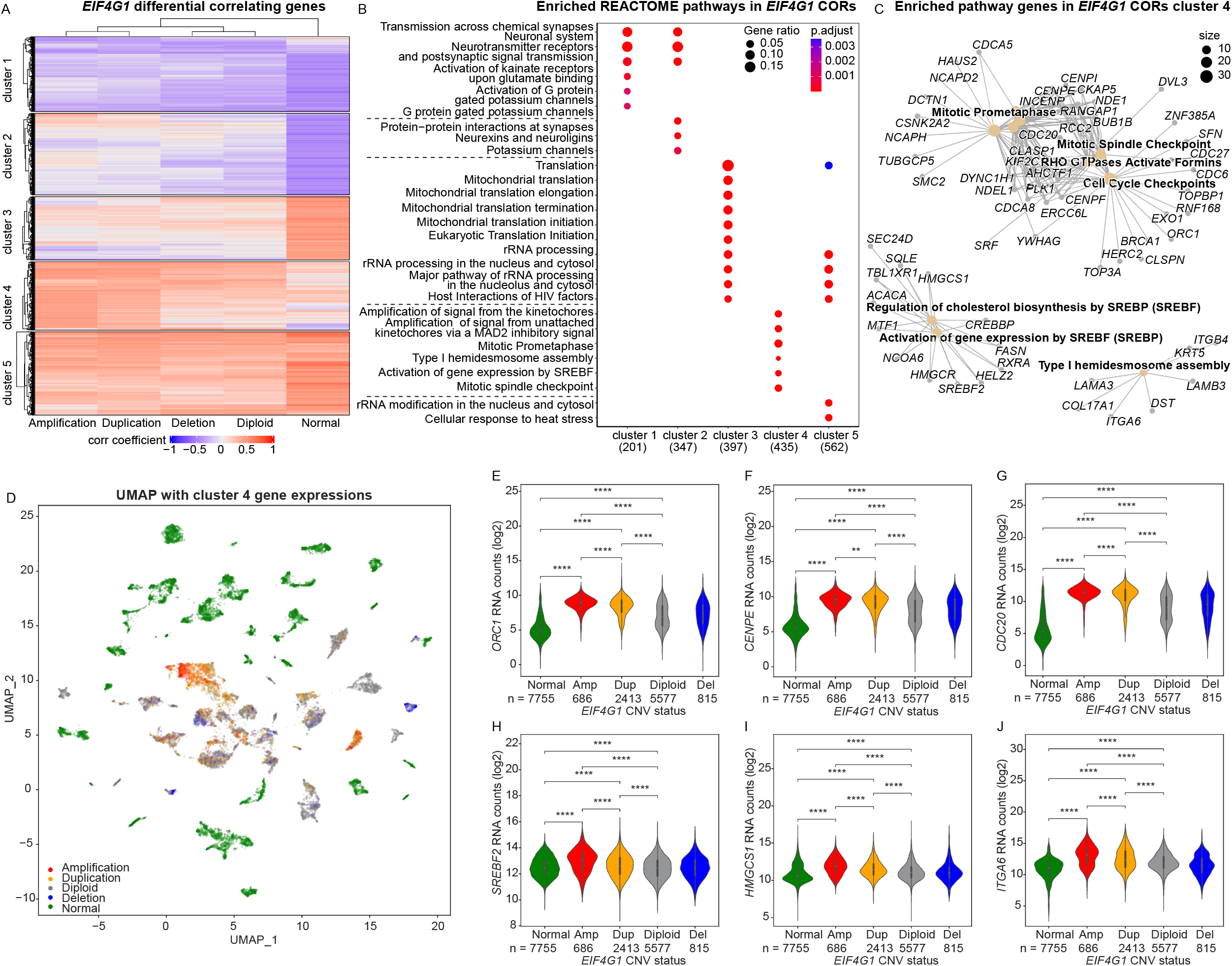
A strong correlation between *EIF4G1* expression and cancer survival genes in tumors with *EIF4G1* amplification or duplication. (*A*) The heatmap illustrates the differential expression correlation between cellular genes and *EIF4G1* in TCGA tumor samples with different *EIF4G1* CNV statuses. Pearson’s correlation coefficients between *EIF4G1* and each of 58,582 other genes were calculated separately across TCGA tumor samples with different *EIF4G1* CNV statuses (labeled as Amplification, Duplication, Diploid or Deletion (heterozygous and homozygous deletion)), or across 7,414 GTEx healthy samples from different tissue types (labeled as Normal). Genes with significant positive (r > 0.3) or negative (r < -0.3) correlations were selected for further analysis. Each row indicates the correlation of a gene with *EIF4G1* in the groups with the indicated *EIF4G1* CNV status. The heatmap cells’ color and intensity correspond to Pearson’s correlation coefficient values. The dendrogram at the top shows the hierarchical relationship between the columns. The rows were ordered and partitioned into 5 non-overlapping subgroups using a K-means clustering algorithm. (*B*) The dot plot displays the enriched pathways identified through REACTOME pathway analysis for the heatmap row clusters in (*A*). The 8 most significantly enriched pathways of each cluster are plotted, ranked by their adjusted p-values. (*C*) The star network plot illustrates the genes that belong to the significantly enriched pathways in cluster 4. (*D*) The UMAP provides a visual grouping of mRNA expression of genes in cluster 4 from (*A*). Normal (healthy) samples are shown in green, while other colors represent tumor samples with different EIF4G1 copy number variation statuses. (*E*-*J*) The box plots compare the median expression of genes from cluster 4 in healthy samples and tumor samples with different *EIF4G1* copy number variation statuses. The two-tailed Student’s t-tests were performed. ns, not significant; *p ≤ 0.05; **p ≤ 0.01; ***p ≤ 0.001; ****p ≤ 0.0001.

Moreover, the cluster 4 genes from *EIF4G1* analysis (Fig. 2*A* and *B*) are involved in the pathways crucial for cancer cell survival, such as regulation of cell cycle, cholesterol synthesis and lipogenesis, and cell adhesion pathways (26-28) (Fig. 2C). In contrast, the genes in clusters 3 and 5 had strong positive correlations with *EIF4G1* in healthy tissues, but not in tumors, and are involved in housekeeping pathways such as translation and ribosomal RNA processing (Fig. S4*A* and *B*). These findings suggest that *EIF4G1* positively influences pathways beneficial for cancer survival, especially in tumors with *EIF4G1* gain. However, in tumors, *EIF4G1*’s influence on housekeeping pathways is weaker compared to healthy tissues. Interestingly, in healthy tissues, cluster 1 and cluster 2 genes exhibit anti-correlations with *EIF4G1* and *EIF3E* expression (Figs. 2*A* and S3*A*), which include genes for neuronal proteins, G proteins, and ion channels.

To verify if the observed differential correlations reflect differential gene expression, we used the dimensionality reduction algorithm, uniform manifold approximation and projection (UMAP) (29), to assess cluster 4 gene expressions in healthy tissues and tumors. The UMAP analysis clearly distinguished healthy tissue samples from tumor samples based on the differential expression of cluster 4 genes. Multiple distinct clusters were observed among healthy tissues (green dots in Fig. 2*D*), likely indicating tissue-specific separation. Most tumors with *EIF4G1* amplification or duplication (red or orange dots in Fig. 2*D*) formed separate groupings from tumors with *EIF4G1* diploid or deletion, indicating differential expression of cluster 4 genes within tumor groups. Additionally, UMAP was employed for the expression of clusters 3 and 5 genes in healthy tissues and tumors, which confirmed the differential expression of clusters 3 and 5 genes between healthy tissue and tumors (Fig. S4*C* and *D*).

Finally, we confirmed that the expressions of *EIF4G1* and several cluster 4 genes were significantly elevated in tumors with *EIF4G1* gain (Fig. S4*E*, Fig. 2*E* to *J*, Fig. S4*S* to *H*), suggesting increased activities of cancer survival genes co-regulated with *EIF4G1*, particularly in tumors with *EIF4G1* gain. Altogether, these findings suggest that *EIF4G1* gain is likely a result of positive selection for its tumor-promoting effects.

### Dysregulated eIF4F complex subunits are essential for cancer cell survival

Co-occurring amplification of specific initiation factors may indicate dysregulation of initiation complexes in cancers. To investigate the dysregulation of eIF4F, we first analyzed the correlations among eIF4G1, eIF4A1, eIF4E, 4E-BP1, eIF3E, and eIF3H protein levels in 375 cell lines from CCLE (30). Strong positive correlations were found between eIF4A1 and eIF4G1 (r = 0.632), eIF4E and 4E-BP1 (r = 0.534), eIF3E and eIF3H (r = 0.885), eIF4G1 and eIF3E (r = 0.634), eIF4G1 and eIF3H (r = 0.563), eIF4A1 and eIF3E (r = 0.679), and eIF4A1 and eIF3H (r = 0.650) (Fig. 3*A*). However, eIF4E showed weak positive correlations with eIF4G1 (r = 0.358) and no correlations with eIF4A1, eIF3E, or eIF3H.

**Figure 3.**
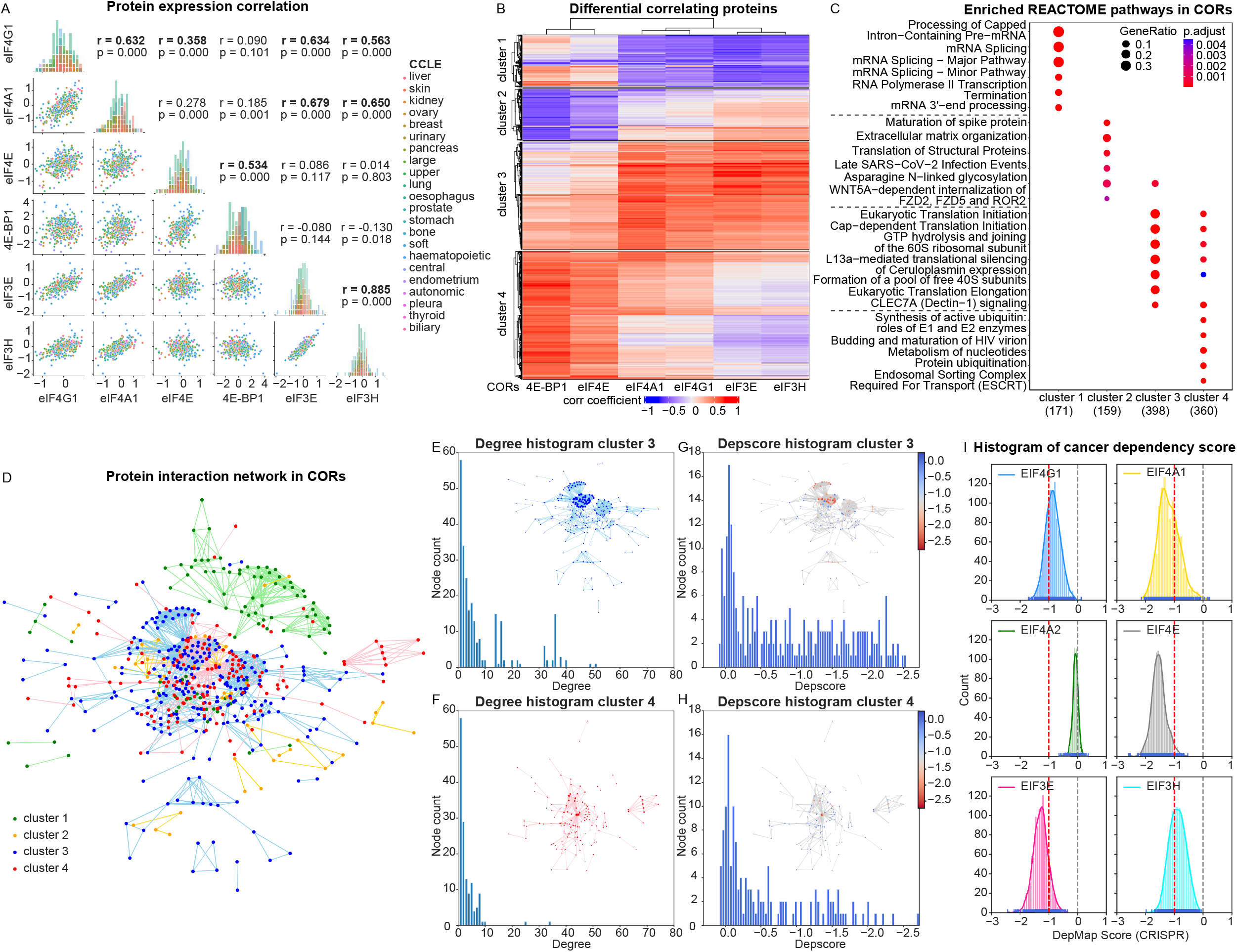
The eIF4F complex components are vital for cancer cell survival and show dysregulation. (*A*) The scatter plots display the correlation between the protein expression levels of eIF4G1, eIF4A1, eIF4E, 4E-BP1, eIF3E, and eIF3H across CCLE cancer cell lines. Bold font denotes strong positive correlations. The colors in the plot represent the tissue origins of the cancer cell lines. Scatterplot axes show the protein expression levels (quantified as the relative abundance of detected peptides to reference, and log2 transformed) from CCLE proteomics data. The histograms show binned expression distribution of each protein (by column) across cancer line cells. For histograms, x-axes match scatterplot x-axes by column, y-axes (unlabeled) depict frequency counts, and coloring matches scatterplots. (*B*) The heatmap displays the differential expression correlation between cellular proteins and eIF4G1, eIF4A1, eIF4E, 4E-BP1, eIF3E, or eIF3H across CCLE cancer cell lines. Pearson’s correlation coefficients between eIF4G1 and each of 12,755 other cellular proteins were calculated separately across 375 CCLE cancer cell lines. Proteins with strong positive (r > 0.5) or negative (r < -0.5) correlations were selected for analysis. (*C*) The dot plot shows the enriched pathways for the heatmap row clusters in (*B*), according to REACTOME pathway analysis. (*D*) The protein-protein interaction network shows the potential interactions between the proteins within clusters identified in the heatmap (*B*). The network was generated using evidence from the STRING database and includes proteins from all clusters. The colors in the network represent the cluster to which each protein (node) belongs, and the potential interactions (edges) between them. (*E* and *F*) The histograms depict the distribution of node degree for the networks built from proteins in clusters 3 and 4 (as shown in the insets). In the insets, the node size reflects its degree -the number of connections it has to other nodes. (*G* and *H*) The histogram displays the distribution of the dependency score (a metric for gene essentiality measured through CRISPR knockout screens in cancer cell lines(50)) for each node in the network. In the insets, the node size represents the degree, and the color reflects their median dependency score across CCLE cancer cell lines. (*I*) The kernel density plots show the distribution of dependency scores for a protein across all CCLE cancer cell lines. A lower score indicates that the gene is more likely to be essential for cancer cell survival in a particular cell line. A score of 0 indicates that the gene is not essential, while a score of -1 represents the median of all commonly essential genes.

We then identified cellular proteins strongly correlated with these proteins, grouped them into four clusters (Fig. 3*B*), and conducted pathway enrichment analysis (Fig. 3*C*). Cluster 3 proteins positively correlate with eIF4G1, eIF4A1, eIF3E, and eIF3H and are involved in translation initiation and ribosomal large and small subunits (Fig. S5*A*). Cluster 4 proteins positively correlate with eIF4E and 4E-BP1, but not with eIF4G1, eIF4A1, eIF3E, or eIF3H. These proteins are involved in ubiquitination, nucleotide metabolism, and endosomal sorting complexes required for transport machinery (ESCRT) pathways (Fig. S5*D*). Cluster 1 proteins negatively correlate with eIF4G1, eIF4A1, eIF3E, and eIF3H and are involved in mRNA splicing (Fig. S6*A*), while cluster 2 proteins negatively correlate with eIF4E and 4E-BP1 and participate in extracellular matrix organization and viral infection pathways (Fig. S7*A*). These findings indicate distinct co-regulation mechanisms for eIF4A1, eIF4G1, eIF3E, and eIF3H compared to eIF4E and 4E-BP1, suggesting dysregulation of cap-dependent initiation and potential cap-independent initiation mechanisms in cancer cells.

To identify the protein complexes involved in co-regulation pathways, we constructed a protein-protein interaction network using the STRING dataset (Fig. 3*D*). Cluster 3 contains numerous subunits from complexes such as the ribosome, eIF2, eIF3, and eIF4F. Our analysis of the cluster 3 network’s degree centrality (Fig. 3*E* and Fig. S5*B*) revealed extensive interactions among its proteins. Additionally, we investigated the relationship between protein connectivity and essentiality by examining the depscores (31) of nodes (Fig. 3*G* and Fig. S5*C*). We found a significant number of high connectivity protein nodes in cluster 3, and these proteins also displayed essential depscores. Within the cluster 4 network, two ubiquitin ribosomal fusion proteins, Uba52 and Rps27A (32), were heavily connected to other proteins (Fig. 3*F* and Fig. S5*E*) and are highly essential based on depscores (Fig. 3*H* and Fig. S5*F*). These findings hint at a co-regulatory relationship between eIF4E activity and ubiquitination in cancer cells. Furthermore, the cluster 1 network exhibited modest connectivity with essential subunits (Fig. S6*B* and *C*), primarily consisting of the mRNA splicing complex (Fig. S6*D* and *E*). In contrast, the cluster 2 network showed low connectivity (Fig. S7*B* and *C*) and lacked essential components (Fig. S7*D* and *E*).

Finally, to compare the essentiality of eIF4F and eIF3 subunits, we plotted the distribution of their depscores across CCLE cancer cell lines (Fig. 3*I*). eIF4G1, eIF4A1, eIF4E, eIF3E, and eIF3H are essential for viability in most cancer cell lines, whereas eIF4A2 is not. In summary, eIF4G1 and eIF4A1 co-regulate with essential ribosome, eIF2, and eIF3 complexes in cancer cell survival. Additionally, eIF4E strongly co-regulates with vital ubiquitin-synthesis complexes. These findings indicate dysregulations of eIF4E and eIF4G1 in cancer cells, suggesting the existence of separate initiation mechanisms: cap-dependent and cap-independent mechanisms, both crucial for cancer cell viability.

### Predictive structure modeling indicates distinct conformations of the eIF4G1 HEAT-2 domain with and without eIF4E binding

Using Alphafold multimer prediction, we modeled the structure of the eIF4F complex in two initiation mechanisms. We modeled the canonical eIF4F complex for cap-dependent initiation, consisting of eIF4G1_557-1437_ (including eIF4E binding domain (4E-BD), HEAT-1, CY, and HEAT-2), along with full-length eIF4A1 and eIF4E (Fig. 4*A*). We modeled a dysregulated conformation of eIF4F for cap-dependent initiation by including only eIF4G1_557-1437_ and full-length eIF4A1. Additionally, we modeled the eIF4E and 4E-BP1 complex (Fig. 4*A*).

**Figure 4.**
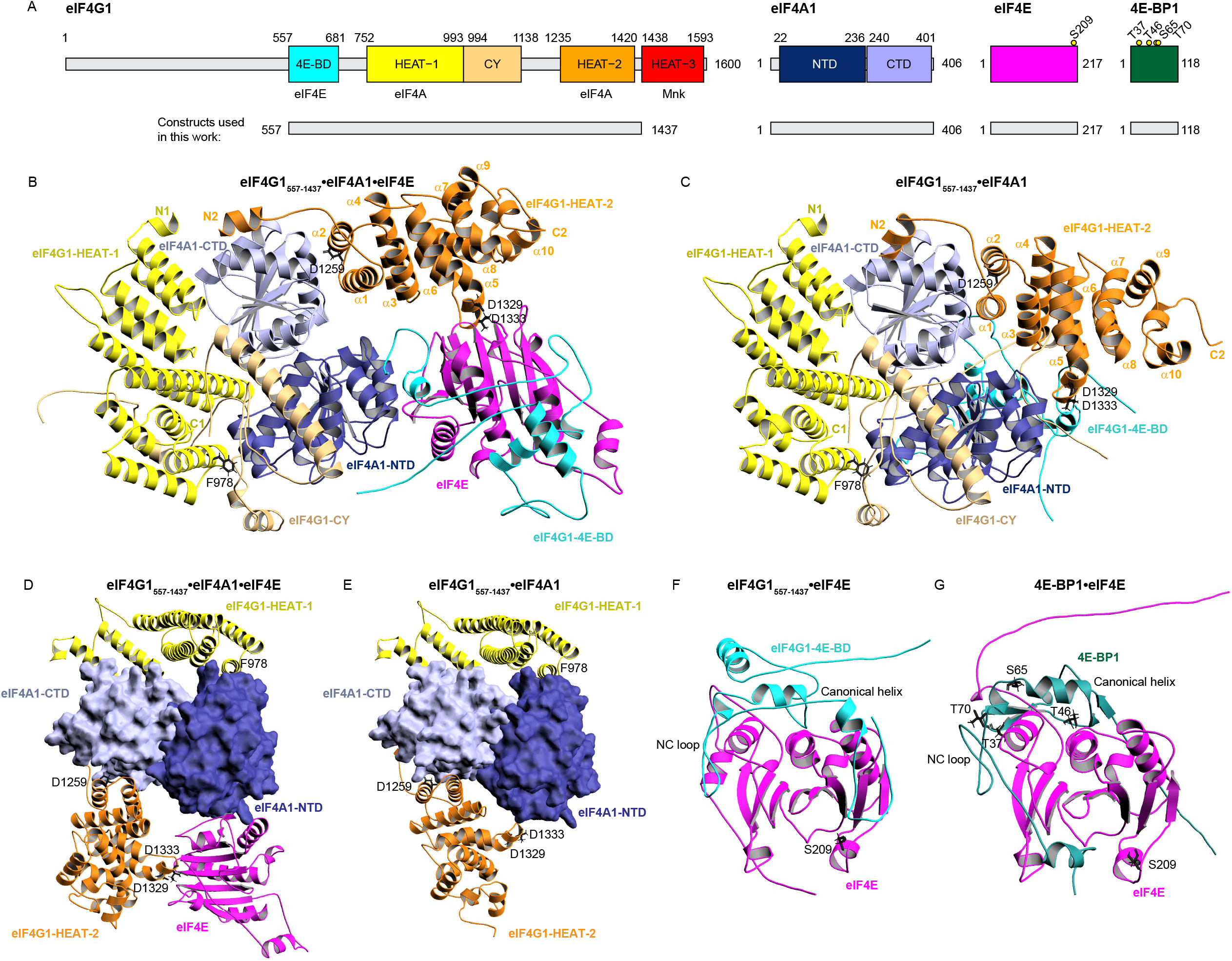
HEAT2 domain of eIF4G1 takes different conformations when eIF4G1 binds to eIF4E (*A*) The diagram illustrates the domain organization of eIF4G1, eIF4A1, eIF4E, and 4E-BP1 constructs used in this work. (*B* and *C*) The predicted protein structures for the eIF4G1•eIF4A1•eIF4E and eIF4G1•eIF4A1 complexes, respectively, as determined by the AlphaFold2 multimer approach. Labeled residues in eIF4G1 have been reported to influence eIF4A binding. (*D* and *E*) the eIF4G1-binding surfaces of eIF4A1 from a different perspective, with eIF4A1 displayed in surface representation and eIF4G1 domains and eIF4E shown in ribbon representation. (*F* and *G*) Comparison of Alphafold-predicted protein structures for eIF4G1(4E-BD) and eIF4E from the eIF4G1_557-1437_•eIF4A1•eIF4E complex in (*B*), and 4E-BP1•eIF4E complexes. eIF4E is shown in surface representation. Labels are given to the residues in eIF4E and 4E-BP1 that are reported phosphorylation sites and could affect their interactions.

In both eIF4G1_557-1437_•eIF4A1•eIF4E (Fig. 4*B* and *D*, Fig. S12*A* and *B*) and eIF4G1_557-1437_•eIF4A1 complexes (Fig. 4*C* and *E*, Fig. S12*C* and *D*), the two eIF4A1-binding domains in eIF4G1 (HEAT-1 and HEAT-2, colored yellow and orange, respectively) are modeled on opposite sides of eIF4A1. HEAT-1 interacts with both the C-terminal domain (CTD, light blue) and N-terminal domain (NTD, dark blue) of eIF4A1. The C-terminal helix of HEAT-1 has been experimentally verified to bind to eIF4A1-NTD (33), with a point mutation in F978 affecting this binding (20). Furthermore, HEAT-2 was predicted to interact with eIF4A1-CTD through a small loop linking α-helix_1_ and α-helix_2_ (33). Indeed, former experimental studies have verified that D1259 is within this loop at the interaction interface between HEAT-2 and eIF4A1-CTD (33).

In the eIF4G1_557-1437_•eIF4A1•eIF4E complex, the Alphafold prediction suggests that 4E-BD binds to eIF4E, bringing eIF4E close to eIF4A1-NTD. The α-helix_5_ of HEAT-2 is positioned near the cap-binding loop of eIF4E, and both HEAT-2 and eIF4E are in proximity to the mRNA binding cavity in eIF4A1. In contrast, in the eIF4G1_557-1437_•eIF4A1 complex, HEAT-2 α-helix_5_ was predicted to directly interact with eIF4A1-NTD via the D1329 and D1333 residues of HEAT-2, a finding supported by experimental studies (22, 33). These findings indicate that in the absence of eIF4G1-eIF4E binding, HEAT-2 interacts with both the CTD and NTD of eIF4A1, resulting in a stronger interaction between eIF4G1 and eIF4A1 in the complex.

To investigate the strong co-regulation between eIF4E and 4E-BP1, we compared the predicted 4E-BD and eIF4E interaction in the eIF4G1_557-1437_•eIF4A1•eIF4E complex (Fig. 4*F*) to the predicted 4E-BP1 and eIF4E interaction in the 4E-BP1•eIF4E complex (Fig. 4*G*). The predicted interactions between the dorsal side of eIF4E and the canonical α-helix binding motifs of 4E-BD or 4E-BP1 are consistent with prior research findings (34-36). The non-canonical loop (NC-loop) from 4E-BD or 4E-BP1 was predicted to bind to the lateral side of eIF4E, as previously reported (34, 36, 37). Phosphorylation of T37, T46, S65, and T70 have been reported to affect 4E-BP1 binding to eIF4E, and those residues were predicted to be at the interaction interface between two proteins. Furthermore, Alphafold prediction places the C-terminal of 4E-BP1 at the cap-binding site of eIF4E, though experimental verification is required.

### Discussion

The frequent amplification of translation initiation genes in TCGA tumor samples highlights their potential importance in tumor function. Our study identified several translation initiation genes including *EIF4G1, EIF3E, EIF3H, EIF4A2, EIF2B5*, and *PABPC1*, as well as several ribosomal large-subunit genes, are co-located in two highly amplified chromosome loci, 3q27 and 8q23, in human cancers. The co-occurrence of copy number gain of *EIF4G1* at 3q27 and *EIF3E* at 8q23 from unlinked loci is noteworthy, as their encoded proteins play an essential role in ribosome recruitment on mRNA (10). Furthermore, we discovered that *PABPC1* proximal to *EIF3E* at 8q23 is also co-gained with *EIF4G1*, and their encoded proteins’ biochemical interaction enhances initiation and ribosome recruitment (38). While functionally related metabolic genes often cluster in close genomic proximity for co-expression and are subject to similar structural variations (39), the co-occurrence of genetic alterations in genetically unlinked driver genes that activate collaborating oncogenic pathways is also commonly observed (40). Our findings suggest positive selections for interactions between eIF4G1 and subunits from other initiation complexes, implying their potential coordinated roles in human cancers.

Positive selection of *EIF4G1* amplification likely results from a feedback mechanism that regulates the expression of cell cycle and lipogenesis genes. Our findings show a strong correlation between *EIF4G1* expression and both cell cycle genes and SREBF-controlled lipogenesis genes in tumors with *EIF4G1* gain. The cell cycle and lipogenesis genes, as identified in Fig. 2, are not genetically linked to *EIF4G1*, yet they are overexpressed in tumors with *EIF4G1* gain. This suggests that cancer clones with *EIF4G1* gain may be positively selected because it facilitates the overexpression of cancer survival genes. Additionally, research has shown that *SREBF1* translation relies on cap-independent initiation under stress conditions that inhibit cap-dependent initiation (41). Further studies are required to evaluate whether inhibition of translation initiation could break the feedback loop and impede tumor growth, through the *SREBF* lipogenesis axis.

The strong co-regulation of eIF4G1, eIF4A1, eIF3E, and eIF3H, along with the independent co-regulation of eIF4E and 4E-BP1, suggests additional regulatory mechanisms for eIF4E beyond its role in the eIF4F complex. The co-regulation of eIF4G1, eIF4A1, eIF3E, and eIF3H with ribosomal subunits and various initiation complexes underscores their role in initiating translation, vital for cancer cell survival (Fig.4).

Interestingly, eIF4E is essential for cancer cell viability, and it also co-regulates with 4E-BP1, ubiquitination, and ESCRT proteins. Previous studies have shown that ubiquitination and 4E-BP1 can reduce eIF4E’s ability to bind to eIF4G1, thus hindering cap-dependent initiation (42). Dysregulation of eIF4E and eIF4G1 in cancer cells could enable cap-independent mechanisms to meet critical translation initiation needs (3). Furthermore, ESCRT’s role in binding to ubiquitinated proteins and its known regulation of the nuclear pore complex (NPC) (43), may facilitate eIF4E’s mRNA export function. Dysregulation in cap-dependent translation initiation may result from an increased allocation of eIF4E to nuclear mRNA export, reducing its availability in the cytoplasmic eIF4F pool. Hence, eIF4E’s essentiality may stem from both its mRNA transport function alongside its crucial role in cap-dependent initiation.

The Alphafold predictions propose potential mechanisms to regulate eIF4A1’s RNA helicase activity involving the α-helix_5_ of HEAT-2, which can interact with either eIF4E or eIF4A1-NTD. eIF4G1’s 4E-BD binds to eIF4E, positioning the cap-binding site of eIF4E close to the mRNA binding cavity of eIF4A1 (44), allowing eIF4E and HEAT-2 to encircle the cavity. In the absence of eIF4E, the interaction between HEAT-2 α-helix_5_ and eIF4A1-NTD might enhance eIF4G1 and eIF4A1 interaction, resulting in a tighter mRNA encirclement. Furthermore, the predicted model places 4E-BD close to the mRNA binding cavity of eIF4A1, and previous studies have shown that eIF4G1’s 4E-BD inhibits eIF4A1 helicase activity in the absence of eIF4E binding (9). Further investigations are needed to understand whether additional eIF4A1 molecules could compensate for mRNA unwinding activity in cap-independent initiation and whether cancer cells relying on cap-independent initiation mechanisms may be more susceptible to targeted eIF4A1 inhibition due to their increased dependence on eIF4A1’s helicase activity.

Finally, our study proposes an intriguing hypothesis: eIF4E is crucial for cap-dependent initiation in human cancers; however, in situations of elevated mRNA transport demands, eIF4E may dissociate from the eIF4F complex. In this scenario, we postulate that the dysregulated eIF4F complex, mainly consisting of eIF4G1 and eIF4A1, could remain on the mRNA, potentially enabling cap-independent initiation via eIF3, ribosome, and mRNA circularization. Nonetheless, further experimental investigations are essential to validate this hypothesis.

## Materials and Methods

### Copy number variation data and co-occurrence analysis

Gene-level copy number data for 33 TCGA cancer types were obtained from the UCSC Xena data hub (45) (https://tcga.xenahubs.net and https://pancanatlas.xenahubs.net). To classify copy number variation statuses, the TCGA pan-cancer gene-level CNV threshold dataset we used, which combined GISTIC2-thresholded data from all TCGA cohorts, accessed through the Xena dataset ID: TCGA.PANCAN.sampleMap/Gistic2_CopyNumber_Gistic2_all_thresholded.by_genes. We grouped the estimated gene-level CNV values using thresholds 2, 1, 0, -1, -2, to represent high-level copy number gain (amplification), low-level copy number gain (duplication), diploid, shallow (possibly heterozygous) deletion, or deep (possibly homozygous) deletion.

To generate Likert plots of CNV statuses across different cancer types, we utilized clinically relevant phenotype information for TCGA samples, such as sample type and primary disease annotations, obtained from individual TCGA cohorts through the Xena dataset ID: TCGA_phenotype_denseDataOnlyDownload.tsv. To conduct the co-occurrence analysis, we employed the VennCounts() function from the R package “limma” to calculate the number of overlapping genes between gene groups. We then utilized these counts to create proportional Venn diagrams using the euler() function from the R package “eulerr”. Statistical analysis was performed using the fisher.test() function from the R package ‘stats’.

Kaplan-Meier analysis was conducted using curated clinical data from the TCGA Pan-Cancer Clinical Data Resource (46), obtained through the Xena dataset ID: Survival_SupplementalTable_S1_20171025_xena_sp. Survival analysis utilized the fit() function from the class KaplanMeierFitter() in the Python package “lifelines”. Differences in survival curves were evaluated using the log-rank test with the logrank_test() function from the statistics() class in the “lifelines” package.

### RNA-Seq and gene expression analysis

The original RNA-Seq data of tumor samples were obtained from TCGA and RNA-Seq data of healthy samples from GTEx (47). To ensure consistency and minimize computational batch effects on read alignment and quantification, we used the reprocessed RNA-Seq read count data for both sources, available from the UC Santa Cruz computational genomics Lab, which was computed with the Toil-based RNA-Seq bioinformatic pipeline (48). We accessed the RNA-Seq datasets from the UCSC Xena data hub (https://toil.xenahubs.net) using the Xena dataset IDs: TcgaTargetGtex_RSEM_hugo_norm_count.

For the differential correlation analysis, we utilized the corrwith() function from the Python package “pandas” to calculate Pearson’s correlation coefficient (r). To perform UMAP, we standardized the gene expression data by scaling it to unit variance using the fit_transform() function from the class StandardScaler() of the Python package “sklearn.preprocessing”. Next, we used the fit_transform() function from the class UMAP() of the Python package “umap” to embed the standardized data into a Euclidean space.

### Proteomics and network analysis

The CCLE proteomics data were obtained from the publication (30). We obtained the dependency score data of CRISPR knockout screens and sample information from the depmap portal, using the file name “CRISPR_gene_effect.csv” and “sample_info.csv”. To construct the protein-protein interaction network, we used protein network data from the STRING database with the file name “9606.protein.physical.links.detailed.v11.5.txt”. We constructed the protein-protein interaction networks using the from_pandas_edgelist() function and plotted them with the draw_networkx() and kamada_kawai_layout() functions from the Python package “networkx”. We conducted the centrality analysis on the network using the degree_centrality() function from “networkx”.

### Heatmap, clustering, and pathway enrichment analysis

To create heatmaps, we utilized the Heatmap() function from the R package “ComplexHeatmap”. The heatmap rows were ordered and grouped into subgroups using the K-means clustering method while the heatmap columns were ordered by the hierarchical clustering method. To analyze the enriched biological pathways of the genes within each cluster, we used the enrichPathway() function from the R package “ReactomePA”, which employed Reactome as a source of pathway data. We performed statistical analysis and visualization of these pathways using the compareCluster() function from the R package “clusterProfiler”, by applying an over-representation analysis (ORA) method. The statistical significance (p-value) of the overlap between genes from a given pathway and the gene list was determined using the hypergeometric distribution test, and the p-values were adjusted for multiple comparison using Hochberg’s and Hommel’s method.

### Protein complex structure prediction

We employed the AlphaFold multimer approach, which is an extension of the AlphaFold2 algorithm and is capable of predicting the structure of a protein complex as a single entity (49). All tasks were performed on the GPU cluster of Harvard medical school, with the default AlphaFold2 multimer settings. We used the amino acid sequences from the NCBI database with the following accession number: eIF4E, NP_001959.1; eIF4G1, NP_886553.3; eIF4A1, NP_001407.1 and 4E-BP1, NP_004086.1. We evaluated the consistency of the five prediction models generated by each task and selected the top-ranked model for illustration using PyMOL.

## Data, Materials, and Software Availability

The detailed source code, which reproduces the results reported in this manuscript, is available online at https://github.com/a3609640/pyEIF in the form of R and Python scripts. All the data utilized in our analyses are publicly accessible and the scripts for downloading them from the relevant data repositories are provided in our software repository.

## Acknowledgments

The results published here are based upon data generated by the TCGA Research Network: https://www.cancer.gov/tcga, the Genotype-Tissue Expression (GTEx) Project, and Cancer Cell Line Encyclopedia (CCLE). This work was supported by the National Cancer Institute (5R01CA200913-05, to G.W.) and the National Institute of Allergy and Infectious Diseases (5P01AI143565-03, to G.W.).

## Notes

### Competing Interest Statement

G.W. is a co-founder of and has equity in Enanta Pharmaceuticals, PIC Therapeutics, Eutropics, Olaris Therapeutics, Skinap Therapeutics, Cellmig Biolabs, NOW Scientific, Virtual Discovery, and QuantumTx.

