## Supporting information for "Computational inference of eIF4F complex function and structure in human cancers"

**This PDF file includes:**

Figures S1 to S8
Tables S1

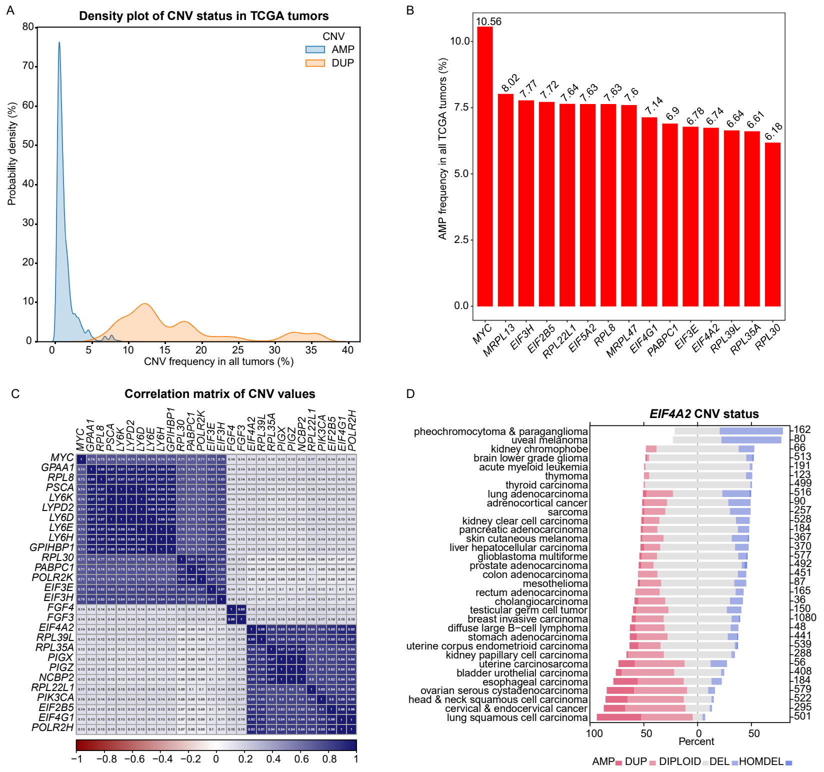


**Fig. S1. Genetic linkage and amplification frequency of translation initiation genes in TCGA tumors**. (*A*) The kernel density estimate plot illustrates the frequency of amplification or duplication of cellular genes in tumors from all cancer types in the TCGA database. (*B*) The bar plot shows the amplification frequency of translation initiation genes, with *MYC*, as a control gene showing the highest amplification frequency among all cellular genes. (*C*) The matrix plot displays Pearson correlation coefficients for estimated CNV values of gene pairs across 10,845 tumors from all TCGA cancer types. The color of each cell reflects the magnitude of the correlation coefficient, and cells with an "X" indicate statistical insignificance ($p > 0.05$, p values not shown). (*D*) The stacked bar plot displays the CNV status for *EIF4A2* genes in different cancer types.

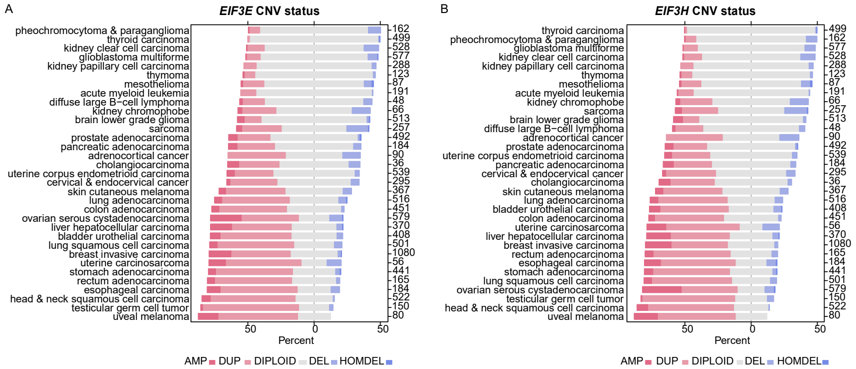


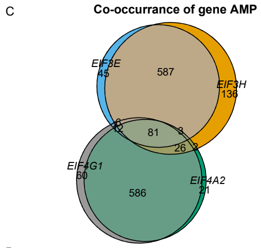


D

| | *EIF3E* AMP | *EIF3E* no AMP | Fisher's exact test (one side) |
|---|---|---|---|
| *EIF4G1* AMP | 100 | 675 | Odd ratios p-value |
| *EIF4G1* no AMP | 635 | 9436 | 2.2045 < 0.001 |

| | *EIF3H* AMP | *EIF3H* no AMP | Fisher's exact test (one side) |
|---|---|---|---|
| *EIF4G1* AMP | 115 | 659 | Odd ratios p-value |
| *EIF4G1* no AMP | 728 | 9343 | 2.23934 < 0.001 |

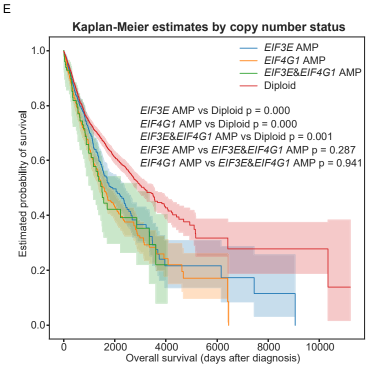


**Fig. S2. Co-amplification of *EIF4G1* and *EIF3E* independent of genetic linkage**. (*A*) The stacked bar plot displays the CNV status for *EIF3E* genes in different cancer types. (*B*) The stacked bar plot displays the CNV status for *EIF3H* genes in different cancer types. (*C*) Venn diagrams show the co-occurrence of amplification in copy number status for the indicated genes among tumors in the TCGA database. (*D*) Co-occurrence statistics. (*E*) The KM plot displays the survival probabilities of TCGA patients with cancer based on the amplification of *EIF4G1* and/or *EIF3E* (indicated inside each box). The statistical significance of differences was determined by p values obtained from log-rank tests. The shaded areas represent a 95% confidence region for the curve.

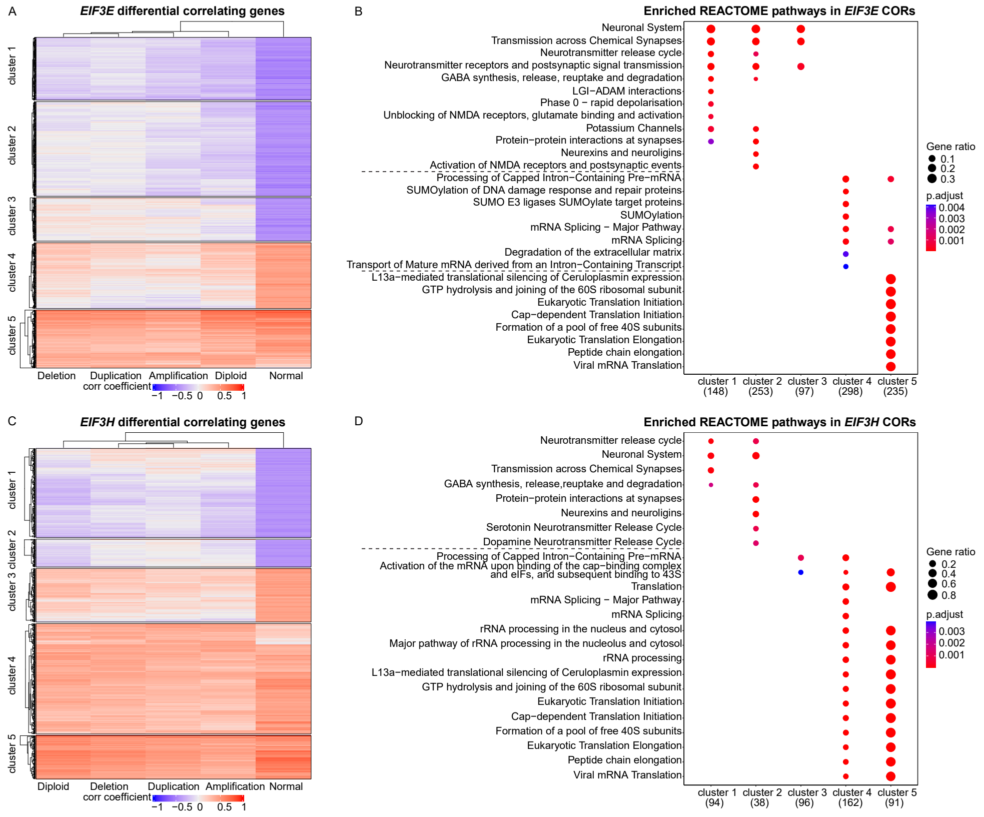


**Fig. S3. mRNA expression-pathway correlations are unaffected by copy number gain for *EIF3E* and *EIF3H*.** (*A*) The heatmap shows the correlation of *EIF3E* mRNA expression with other cellular genes in TCGA tumor samples stratified by *EIF3E* CNV status and GTEx healthy tissues. Pearson's correlation coefficients for *EIF3E* versus 58,582 other genes were calculated separately across TCGA tumor samples with different *EIF3E* CNV statuses (labeled as Amplification, Duplication, Diploid, or Deletion (heterozygous and homozygous deletion)), or across 7,414 GTEx healthy samples from different tissue types (labeled as normal). Genes with significant positive (r > 0.3) or negative (r < -0.3) correlations were included. Each row indicates the correlation of a gene with *EIF3E* in the groups with the indicated *EIF3E* CNV status. The dendrogram at the top indicates similarity as a hierarchical relationship between the columns. The rows are ordered and partitioned into 5 non-overlapping subgroups using a K-means clustering algorithm. (*B*) The dot plot shows the enriched pathways for the heatmap row clusters in (*A*), according to REACTOME pathway analysis. The 8 most significantly enriched pathways of each cluster were plotted. (*C*) The heatmap shows the correlation of *EIF3H* mRNA expression with cellular genes in TCGA tumor samples with different *EIF3H* copy number variation (CNV) status. (*D*) The dot plot shows the enriched pathways for the heatmap row clusters in (*C*), according to REACTOME pathway analysis.

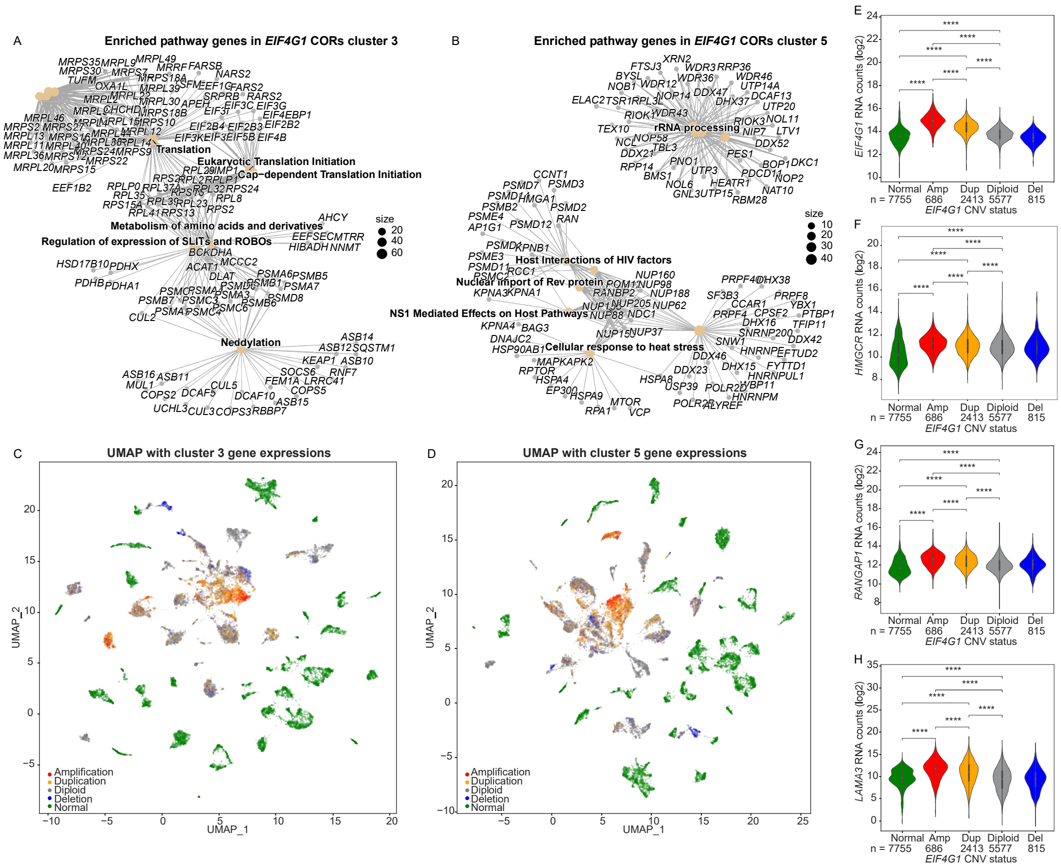


**Fig. S4. Differential correlations of *EIF4G1* with translation-related genes in healthy and tumor samples.** (*A-B*) The star network plots illustrate the genes significantly enriched in the pathways identified in clusters 3 and 5 from Fig. 2*A*. (*C-D*) UMAP visualization of mRNA expression of genes in clusters 3 and 5 from Fig. 2A, where green color indicates normal (healthy) samples, and other colors represent tumor samples with varying CNV statuses. (*E-J*) The box plots compare the median expression of genes in cluster 4 from tumor samples with different *EIF4G1* CNV statuses. The two-tailed Student's t-tests were performed. ns, not significant; ∗p ≤ 0.05; ∗∗p ≤ 0.01; ∗∗∗p ≤ 0.001; ∗∗∗∗p ≤ 0.0001.

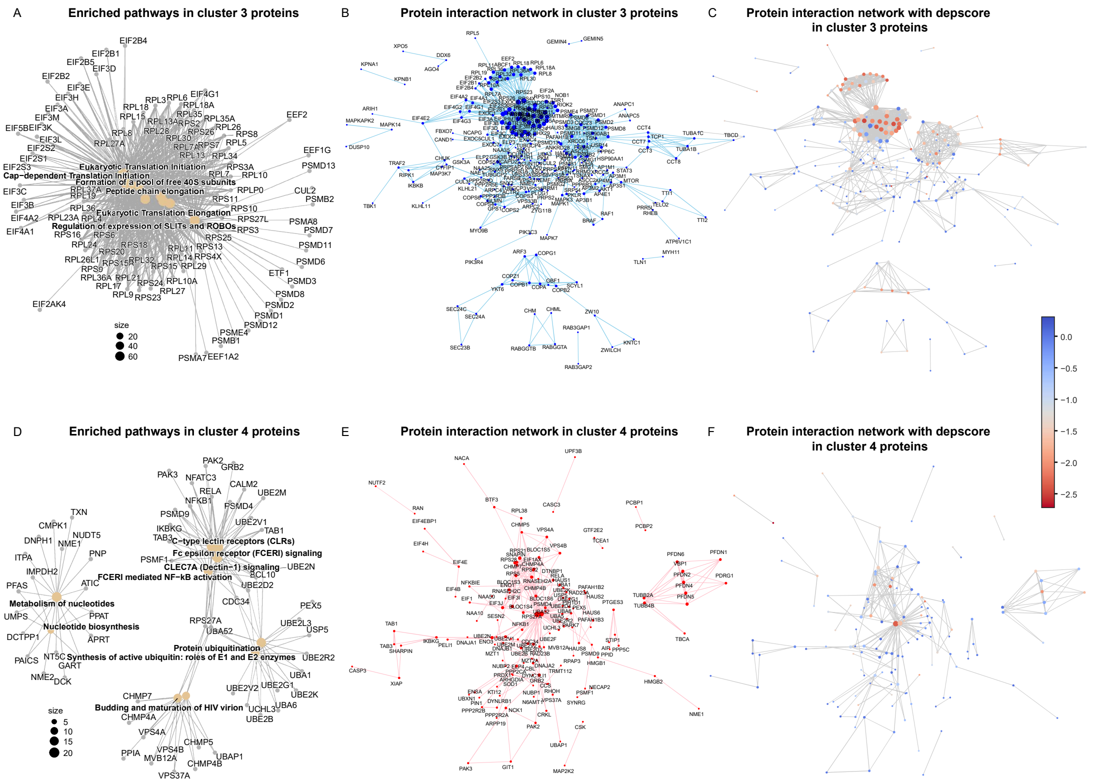


**Fig. S5. Distinct connectivity and dependency score profiles in the protein-protein interaction networks of eIF4G1/eIF4A1 and eIF4E/4E-BP1 co-expression proteins**. (*A*) The star network plot illustrates the genes significantly enriched in the pathways identified in cluster 3 from Fig. 3*B*. (*B*) The enlarged network graph from the inset of Fig. 1*E*. Node size represents the degree, i.e., the number of connections each node has to other nodes. (*C*) The enlarged network graph from the inset of Fig. 1*G*. Node size represents the degree, and node color reflects the median dependency score. (*D*) The star network plot illustrates the genes significantly enriched in the pathways identified in cluster 4 from Fig. 3*B*. (*E*) The enlarged network graph from the inset of Fig. 1*F*. Node size reflects its degree. (*F*) The enlarged network graph from the inset of Fig. 1*H*. Node size represents its degree, and the color reflects its median dependency score.

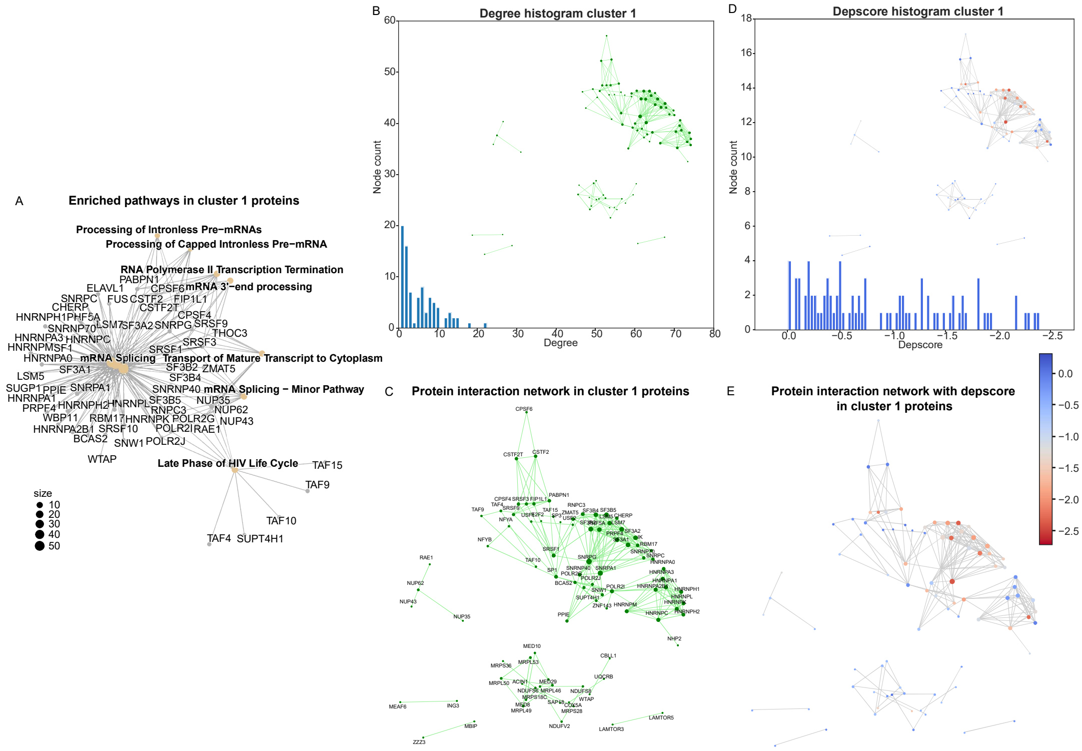

**Fig. S6.** (*A*) The star network plot illustrates the genes that belong to the significantly enriched pathways identified in cluster 1 from Fig. 3*B*. (*B*) The histograms depict the distribution of node degree for the networks built from proteins in cluster 1 (as shown in the insets). Node size represents its degree. (*C*) The enlarged network graph from the inset of (*B*). Node size represents its degree. (*D*) The histogram shows the distribution of the dependency score of each node in the network. (*E*) In the inset, node size represents its degree, and the color reflects its median dependency score.

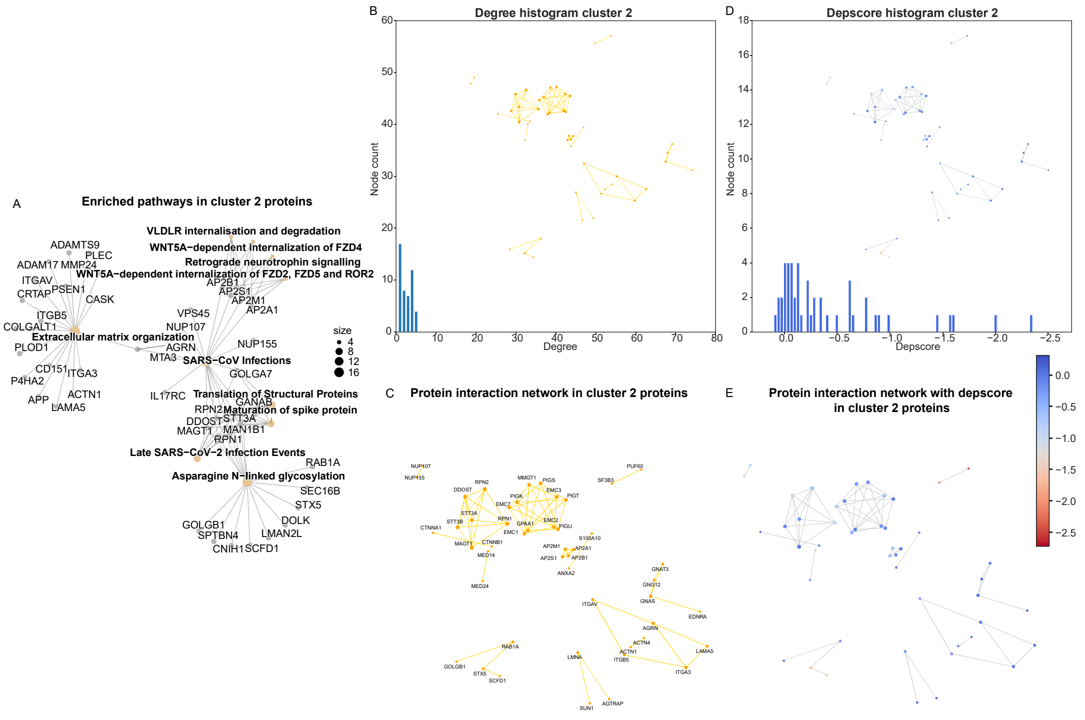


**Fig. S7.** (*A*) The star network plot illustrates the genes that belong to the significantly enriched pathways identified in cluster 2 from Fig. 3*B*. (*B*) The histograms depict the distribution of node degree for the networks built from proteins in cluster 2 (as shown in the insets). The size of each node reflects its degree - the number of connections it has to other nodes. (*C*) The enlarged network graph from the inset of (*B*). The size of each node reflects its degree. (*D*) The histogram shows the distribution of the dependency score of each node in the network. (*E*) In the inset, the size of the nodes represents their degree, and the color reflects their median dependency score.

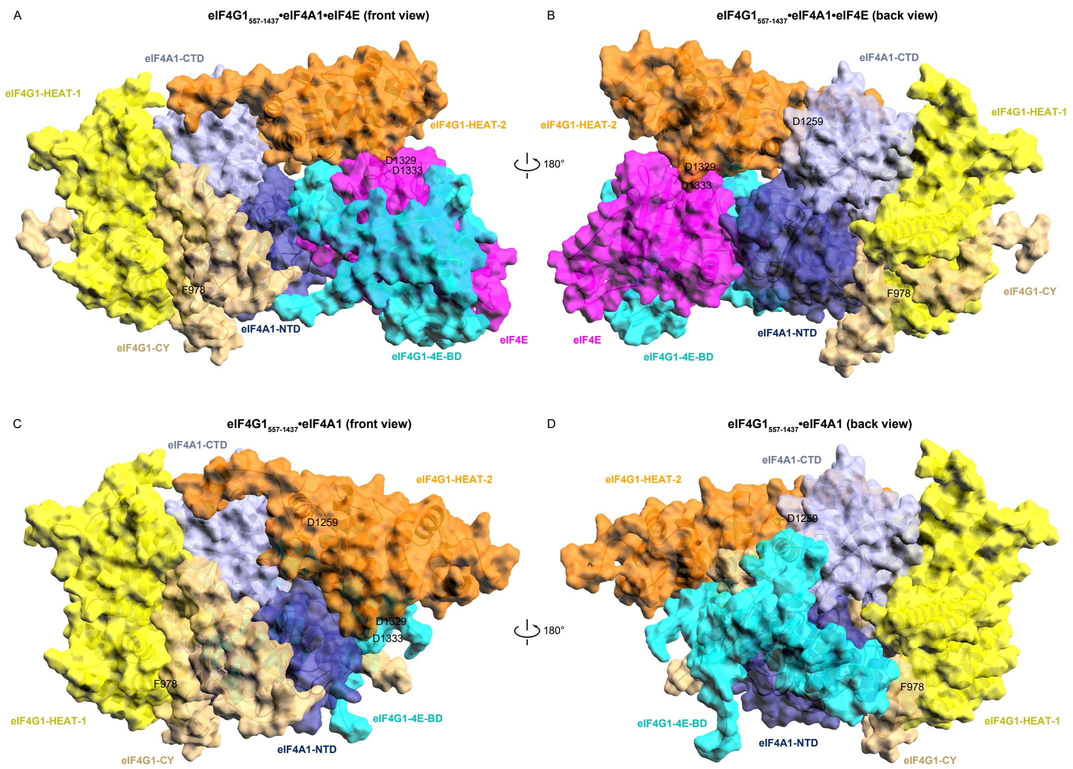


**Fig. S8.** (*A* and *B*) Surface and cartoon representations of the structure of eIF4G1•eIF4A1•eIF4E in two orientations. (*C* and *D*) Surface and cartoon representations of the structure of eIF4G1•eIF4A1 in two orientations.

**Table S1.** Type or paste table title here. Paste table below the title.

| | Gene | AMP Freq % | entrez |
|---|---|---|---|
| 1 | 7SK\|ENSG00000254144.2 | 7.745504841 | |
| 2 | AARD | 7.699400645 | 441376 |
| 3 | ABCC5 | 7.496542185 | 10057 |
| 4 | ABCF3 | 7.210696173 | 55324 |
| 5 | ABRA | 6.740433379 | 137735 |
| 6 | ACAP2 | 6.83264177 | 23527 |
| 7 | ACTL6A | 7.607192254 | 86 |
| 8 | ACTRT3 | 8.114338405 | 84517 |
| 9 | ADAMTSL4 | 5.283540802 | 54507 |
| 10 | ADCK5 | 7.634854772 | 203054 |
| 11 | ADCY8 | 8.658367911 | 114 |
| 12 | ADIPOQ | 6.721991701 | 9370 |
| 13 | AGO2 | 8.068234209 | 27161 |
| 14 | AHRR | 5.265099124 | 57491 |
| 15 | AHSG | 6.73121254 | 197 |
| 16 | ALG3 | 7.136929461 | 10195 |
| 17 | ANGPT1 | 6.851083449 | 284 |
| 18 | ANKRD18DP | 6.611341632 | 348840 |
| 19 | ANKRD46 | 7.035500231 | 157567 |
| 20 | ANO1 | 5.956662056 | 55107 |
| 21 | ANXA13 | 8.57538036 | 312 |
| 22 | ANXA9 | 5.172890733 | 8416 |
| 23 | AP2M1 | 7.496542185 | 1173 |
| 24 | APOD | 6.804979253 | 347 |
| 25 | ARC | 7.588750576 | 23237 |
| 26 | ARHGAP39 | 7.634854772 | 80728 |
| 27 | ARL14 | 5.182111572 | 80117 |
| 28 | ARNT | 5.320424159 | 405 |
| 29 | ASAP1 | 8.944213923 | 50807 |
| 30 | ATAD2 | 8.593822038 | 29028 |
| 31 | ATP11B | 7.607192254 | 23200 |
| 32 | ATP13A3 | 6.685108345 | 79572 |
| 33 | ATP13A4 | 6.657445828 | 84239 |
| 34 | ATP13A5 | 6.611341632 | 344905 |
| 35 | ATP6V1C1 | 6.804979253 | 528 |
| 36 | AZIN1 | 6.795758414 | 51582 |
| 37 | B3GALNT1 | 5.209774089 | 8706 |
| 38 | B3GNT5 | 7.524204703 | 84002 |
| 39 | BAALC | 6.823420931 | 79870 |
| 40 | BAI1 | 7.65329645 | |
| 41 | BCHE | 6.519133241 | 590 |
| 42 | BCL6 | 6.62978331 | 604 |
| 43 | BDH1 | 6.860304288 | 622 |
| 44 | BNIPL | 5.145228216 | 149428 |
| 45 | BOP1 | 7.634854772 | 23246 |
| 46 | BRD9 | 5.265099124 | 65980 |
| 47 | C1orf138 | 5.274319963 | |
| 48 | C1orf56 | 5.145228216 | 54964 |
| 49 | C3orf65 | 6.980175196 | |
| 50 | C3orf70 | 7.090825265 | 285382 |
| 51 | C3orf80 | 5.043798986 | 401097 |
| 52 | C5orf55 | 5.265099124 | |
| 53 | C8orf17 | 7.680958967 | 100507249 |
| 54 | C8orf31 | 7.662517289 | |
| 55 | C8orf33 | 7.634854772 | 65265 |
| 56 | C8orf37 | 5.643153527 | |
| 57 | C8orf47 | 6.150299677 | |
| 58 | C8orf56 | 6.795758414 | |
| 59 | C8orf76 | 8.47395113 | 84933 |
| 60 | C8orf82 | 7.634854772 | 414919 |
| 61 | C8orf87 | 5.163669894 | |
| 62 | CAMK2N2 | 7.127708621 | 94032 |
| 63 | CASC7 | 7.874596588 | |
| 64 | CASC8 | 10.52097741 | 727677 |
| 65 | CCAT1 | 10.0230521 | 100507056 |
| 66 | CCDC127 | 5.265099124 | 133957 |
| 67 | CCDC166 | 7.634854772 | 100130274 |
| 68 | CCDC26 | 9.100968188 | 137196 |
| 69 | CCDC39 | 7.496542185 | 339829 |
| 70 | CCDC50 | 6.556016598 | 152137 |
| 71 | CCND1 | 6.233287229 | 595 |
| 72 | CCNE2 | 5.569386814 | 9134 |
| 73 | CDC42SE1 | 5.163669894 | 56882 |
| 74 | CDH17 | 5.41263255 | 1015 |
| 75 | CEP19 | 6.804979253 | 84984 |
| 76 | CEP72 | 5.265099124 | 55722 |
| 77 | CERS2 | 5.20055325 | 29956 |
| 78 | CGN | 5.062240664 | 57530 |
| 79 | CHRAC1 | 7.865375749 | 54108 |
| 80 | CHRD | 7.173812817 | 8646 |
| 81 | CLCN2 | 7.136929461 | 1181 |
| 82 | CLDN1 | 6.537574919 | 9076 |
| 83 | CLDN11 | 7.985246657 | 5010 |
| 84 | CLDN16 | 6.574458276 | 10686 |
| 85 | CLPTM1L | 5.025357308 | 81037 |
| 86 | COL14A1 | 8.26187183 | 7373 |
| 87 | COL22A1 | 7.75472568 | 169044 |
| 88 | COLEC10 | 8.049792531 | 10584 |
| 89 | COMMD5 | 7.634854772 | 28991 |
| 90 | COX6C | 6.602120793 | 1345 |
| 91 | CPN2 | 6.648224988 | 1370 |
| 92 | CPQ | 5.9197787 | 10404 |
| 93 | CPSF1 | 7.634854772 | 29894 |
| 94 | CRYGS | 6.795758414 | 1427 |
| 95 | CSMD3 | 7.524204703 | 114788 |
| 96 | CT64 | 5.781466113 | |
| 97 | CTHRC1 | 6.694329184 | 115908 |
| 98 | CTSK | 5.255878285 | 1513 |
| 99 | CTSS | 5.348086676 | 1520 |
| 100 | CTTN | 5.698478562 | 2017 |

| | Gene | AMP Freq % | entrez |
|---|---|---|---|
| 101 | CYC1 | 7.736284002 | 1537 |
| 102 | CYHR1 | 7.634854772 | |
| 103 | CYP11B1 | 7.588750576 | 1584 |
| 104 | CYP11B2 | 7.625633933 | 1585 |
| 105 | CYP2AB1P | 7.496542185 | 647265 |
| 106 | DCAF13 | 6.666666667 | 25879 |
| 107 | DCSTAMP | 6.620562471 | 81501 |
| 108 | DCUN1D1 | 7.55186722 | 54165 |
| 109 | DENND3 | 8.086675888 | 22898 |
| 110 | DEPTOR | 8.215767635 | 64798 |
| 111 | DERL1 | 8.307976026 | 79139 |
| 112 | DGAT1 | 7.634854772 | 8694 |
| 113 | DGKG | 6.934071 | 1608 |
| 114 | DLG1 | 6.906408483 | 1739 |
| 115 | DNAJB11 | 6.814200092 | 51726 |
| 116 | DNAJC19 | 7.413554633 | 131118 |
| 117 | DPY19L4 | 5.606270171 | 286148 |
| 118 | DPYS | 6.675887506 | 1807 |
| 119 | DSCC1 | 7.985246657 | 79075 |
| 120 | DVL3 | 7.496542185 | 1857 |
| 121 | EBAG9 | 6.860304288 | 9166 |
| 122 | ECE2 | 7.1461503 | 9718 |
| 123 | ECM1 | 5.117565698 | 1893 |
| 124 | ECT2 | 7.671738128 | 1894 |
| 125 | EEF1D | 7.736284002 | 1936 |
| 126 | EFR3A | 8.446288612 | 23167 |
| 127 | EGFEM1P | 7.85615491 | 93556 |
| 128 | EGFR | 5.053019825 | 1956 |
| 129 | EHHADH | 7.081604426 | 1962 |
| 130 | EIF2B5 | 7.717842324 | 8893 |
| 131 | EIF3E | 6.777316736 | 3646 |
| 132 | EIF3H | 7.773167358 | 8667 |
| 133 | EIF4A2 | 6.740433379 | 1974 |
| 134 | EIF4G1 | 7.136929461 | 1981 |
| 135 | EIF5A2 | 7.634854772 | 56648 |
| 136 | EMC2 | 6.694329184 | 9694 |
| 137 | ENPP2 | 7.948363301 | 5168 |
| 138 | ENSA | 5.357307515 | 2029 |
| 139 | ENY2 | 6.804979253 | 56943 |
| 140 | EPHB3 | 7.238358691 | 2049 |
| 141 | EPPK1 | 7.75472568 | 83481 |
| 142 | ESRP1 | 5.624711849 | 54845 |
| 143 | ETV5 | 6.934071 | 2119 |
| 144 | EXOC3 | 5.265099124 | 11336 |
| 145 | EXOSC4 | 7.773167358 | 54512 |
| 146 | EXT1 | 7.95758414 | 2131 |
| 147 | FADD | 5.781466113 | 8772 |
| 148 | FAM131A | 7.136929461 | 131408 |
| 149 | FAM135B | 7.717842324 | 51059 |
| 150 | FAM157A | 6.611341632 | 728262 |
| 151 | FAM203A | 7.736284002 | |
| 152 | FAM203B | 7.634854772 | |
| 153 | FAM43A | 6.602120793 | 131583 |
| 154 | FAM49B | 8.925772245 | |
| 155 | FAM63A | 5.172890733 | |
| 156 | FAM83A | 8.437067773 | 84985 |
| 157 | FAM83H | 7.634854772 | 286077 |
| 158 | FAM84B | 9.359151683 | |
| 159 | FAM91A1 | 8.584601199 | 157769 |
| 160 | FAM92A1 | 5.237436607 | |
| 161 | FBXL6 | 7.634854772 | 26233 |
| 162 | FBXO32 | 8.547717842 | 114907 |
| 163 | FBXO43 | 6.740433379 | 286151 |
| 164 | FBXO45 | 6.685108345 | 200933 |
| 165 | FER1L6 | 8.639926233 | 654463 |
| 166 | FETUB | 6.721991701 | 26998 |
| 167 | FGF12 | 6.703550023 | 2257 |
| 168 | FGF19 | 6.131857999 | 9965 |
| 169 | FGF3 | 6.011987091 | 2248 |
| 170 | FGF4 | 6.030428769 | 2249 |
| 171 | FNDC3B | 7.893038266 | 64778 |
| 172 | FOXH1 | 7.634854772 | 8928 |
| 173 | FSBP | 5.449515906 | 100861412 |
| 174 | FXR1 | 7.496542185 | 8087 |
| 175 | FYTTD1 | 6.685108345 | 84248 |
| 176 | FZD6 | 6.749654219 | 8323 |
| 177 | GABPB2 | 5.163669894 | 126626 |
| 178 | GDF6 | 5.633932688 | 392255 |
| 179 | GEM | 5.384970032 | 2669 |
| 180 | GHSR | 7.773167358 | 2693 |
| 181 | GLI4 | 7.699400645 | 2738 |
| 182 | GML | 7.634854772 | 2765 |
| 183 | GMNC | 6.509912402 | 647309 |
| 184 | GNB4 | 7.680958967 | 59345 |
| 185 | GOLIM4 | 7.127708621 | 27333 |
| 186 | GOLPH3L | 5.366528354 | 55204 |
| 187 | GP5 | 6.648224988 | 2814 |
| 188 | GPAA1 | 7.773167358 | 8733 |
| 189 | GPIHBP1 | 7.671738128 | 338328 |
| 190 | GPR160 | 7.985246657 | 26996 |
| 191 | GPR20 | 7.745504841 | 2843 |
| 192 | GPT | 7.634854772 | 2875 |
| 193 | GRHL2 | 7.035500231 | 79977 |
| 194 | GRINA | 7.736284002 | 2907 |
| 195 | GSDMC | 8.925772245 | 56169 |
| 196 | GSDMD | 7.745504841 | 79792 |
| 197 | HAS2 | 8.178884278 | 3037 |
| 198 | HES1 | 6.721991701 | 3280 |
| 199 | HHLA1 | 8.390963578 | 10086 |
| 200 | HORMAD1 | 5.338865837 | 84072 |

| | Gene | AMP Freq % | entrez |
|---|---|---|---|
| 201 | HPYR1 | 8.234209313 | 93668 |
| 202 | HRASLS | 6.574458276 | |
| 203 | HRG | 6.749654219 | 3273 |
| 204 | HRSP12 | 6.141078838 | |
| 205 | HSF1 | 7.634854772 | 3297 |
| 206 | HTR3C | 7.496542185 | 170572 |
| 207 | HTR3D | 7.496542185 | 200909 |
| 208 | HTR3E | 7.496542185 | 285242 |
| 209 | IFT80 | 5.080682342 | 57560 |
| 210 | IGF2BP2 | 7.035500231 | 10644 |
| 211 | IL12A | 5.062240664 | 3592 |
| 212 | IL1RAP | 6.556016598 | 3556 |
| 213 | INTS8 | 5.587828492 | 55656 |
| 214 | IQCG | 6.611341632 | 84223 |
| 215 | IQCJ | 5.053019825 | 654502 |
| 216 | JRK | 7.597971415 | 8629 |
| 217 | KCNK9 | 7.763946519 | 51305 |
| 218 | KCNMB2 | 7.727063163 | 10242 |
| 219 | KCNMB3 | 7.745504841 | 27094 |
| 220 | KCNQ3 | 8.418626095 | 3786 |
| 221 | KCNS2 | 6.141078838 | 3788 |
| 222 | KCNV1 | 6.768095897 | 27012 |
| 223 | KHDRBS3 | 8.077455048 | 10656 |
| 224 | KIAA0196 | 8.842784693 | |
| 225 | KIAA0226 | 6.777316736 | |
| 226 | KIAA1429 | 5.486399262 | |
| 227 | KIAA1875 | 7.736284002 | |
| 228 | KIFC2 | 7.634854772 | 90990 |
| 229 | KLF10 | 6.62978331 | 7071 |
| 230 | KLHL24 | 7.496542185 | 54800 |
| 231 | KLHL38 | 8.547717842 | 340359 |
| 232 | KLHL6 | 7.496542185 | 89857 |
| 233 | KNG1 | 6.777316736 | 3827 |
| 234 | KPNA4 | 5.154449055 | 3840 |
| 235 | KRT8P12 | 5.126786538 | 90133 |
| 236 | LAMP3 | 7.496542185 | 27074 |
| 237 | LAPTM4B | 6.177962195 | 55353 |
| 238 | LEPREL1 | 6.768095897 | |
| 239 | LINC00051 | 7.745504841 | 619434 |
| 240 | LINC00501 | 7.588750576 | 1E+08 |
| 241 | LINC00535 | 5.31120332 | |
| 242 | LINC00536 | 7.846934071 | 1E+08 |
| 243 | LINC00568 | 5.117565698 | |
| 244 | LINC00578 | 7.727063163 | 1E+08 |
| 245 | LINC00861 | 9.18395574 | 1E+08 |
| 246 | LINC00884 | 6.657445828 | |
| 247 | LINC00885 | 6.749654219 | 401109 |
| 248 | LINC00887 | 6.666666667 | 1E+08 |
| 249 | LINC00888 | 7.496542185 | 1E+08 |
| 250 | LINC00964 | 8.824343015 | 157381 |
| 251 | LINC00969 | 6.804979253 | |
| 252 | LINC00977 | 9.027201475 | 728724 |
| 253 | LIPH | 7.063162748 | 200879 |
| 254 | LMLN | 6.611341632 | 89782 |
| 255 | LPP | 6.961733518 | 4026 |
| 256 | LRCH3 | 6.666666667 | 84859 |
| 257 | LRP12 | 6.694329184 | 29967 |
| 258 | LRRC14 | 7.634854772 | 9684 |
| 259 | LRRC14B | 5.265099124 | 389257 |
| 260 | LRRC15 | 6.657445828 | 131578 |
| 261 | LRRC24 | 7.634854772 | 441381 |
| 262 | LRRC31 | 8.068234209 | 79782 |
| 263 | LRRC34 | 8.095896727 | 151827 |
| 264 | LRRC6 | 8.26187183 | |
| 265 | LRRIQ4 | 8.086675888 | 344657 |
| 266 | LSG1 | 6.62978331 | 55341 |
| 267 | LY6D | 7.570308898 | 8581 |
| 268 | LY6E | 7.634854772 | 4061 |
| 269 | LY6H | 7.690179806 | 4062 |
| 270 | LY6K | 7.625633933 | 54742 |
| 271 | LYNX1 | 7.570308898 | 66004 |
| 272 | LYPD2 | 7.579529737 | 137797 |
| 273 | LYSMD1 | 5.09912402 | 388695 |
| 274 | MAF1 | 7.736284002 | 84232 |
| 275 | MAFA | 7.727063163 | 389692 |
| 276 | MAGEF1 | 7.24757953 | 64110 |
| 277 | MAL2 | 8.003688336 | 114569 |
| 278 | MAP3K13 | 7.090825265 | 9175 |
| 279 | MAP6D1 | 7.496542185 | 79929 |
| 280 | MAPK15 | 7.634854772 | 225689 |
| 281 | MASP1 | 6.639004149 | 5648 |
| 282 | MATN2 | 6.214845551 | 4147 |
| 283 | MB21D2 | 6.592899954 | 151963 |
| 284 | MCCC1 | 7.561088059 | 56922 |
| 285 | MCF2L2 | 7.607192254 | 23101 |
| 286 | MCL1 | 5.292761641 | 4170 |
| 287 | MECOM | 8.446288612 | 2122 |
| 288 | MED30 | 7.75472568 | 90390 |
| 289 | MFI2 | 6.804979253 | |
| 290 | MFN1 | 7.690179806 | 55669 |
| 291 | MFSD3 | 7.634854772 | 113655 |
| 292 | MIR1205 | 10.00461042 | 1E+08 |
| 293 | MIR1207 | 9.940064546 | 1E+08 |
| 294 | MIR1208 | 9.912402029 | 1E+08 |
| 295 | MIR1224 | 7.136929461 | 1E+08 |
| 296 | MIR1234 | 7.634854772 | 1E+08 |
| 297 | MIR1263 | 5.928999539 | 1E+08 |
| 298 | MIR1273A | 6.657445828 | |
| 299 | MIR151A | 7.976025818 | 442893 |
| 300 | MIR15B | 5.034578147 | 406949 |

| | Gene | AMP Freq % | entrez |
|---|---|---|---|
| 301 | MIR2053 | 7.090825265 | 100302225 |
| 302 | MIR28 | 6.556016598 | 407020 |
| 303 | MIR30B | 8.095896727 | 407030 |
| 304 | MIR30D | 8.105117566 | 407033 |
| 305 | MIR3137 | 6.583679115 | 100422926 |
| 306 | MIR3150B | 5.61549101 | 100500907 |
| 307 | MIR3151 | 6.786537575 | 100422992 |
| 308 | MIR3610 | 7.680958967 | 100500914 |
| 309 | MIR3686 | 8.925772245 | 100500839 |
| 310 | MIR378D2 | 5.292761641 | 100616169 |
| 311 | MIR3919 | 5.016136468 | 100500803 |
| 312 | MIR4257 | 5.265099124 | 100422997 |
| 313 | MIR4448 | 7.496542185 | 100616127 |
| 314 | MIR4456 | 5.265099124 | 100616381 |
| 315 | MIR4457 | 5.080682342 | 100616235 |
| 316 | MIR4471 | 6.915629322 | 100616451 |
| 317 | MIR4635 | 5.375749193 | 100616479 |
| 318 | MIR4662B | 8.769017981 | 100616255 |
| 319 | MIR4663 | 8.427846934 | 100616260 |
| 320 | MIR4664 | 7.634854772 | 100616318 |
| 321 | MIR4789 | 7.533425542 | 100616395 |
| 322 | MIR4797 | 6.795758414 | 100616216 |
| 323 | MIR5194 | 8.833563854 | 100847051 |
| 324 | MIR548A3 | 6.574458276 | 693127 |
| 325 | MIR548AA1 | 8.529276164 | 100500863 |
| 326 | MIR548AQ | 6.998616874 | 100847078 |
| 327 | MIR548K | 5.781466113 | 100313770 |
| 328 | MIR551B | 7.395112955 | 693136 |
| 329 | MIR5680 | 6.685108345 | 100847001 |
| 330 | MIR569 | 7.662517289 | 693154 |
| 331 | MIR570 | 6.804979253 | 693155 |
| 332 | MIR599 | 6.316274781 | 693184 |
| 333 | MIR661 | 7.727063163 | 724031 |
| 334 | MIR720 | 6.011987091 | |
| 335 | MIR875 | 6.316274781 | 100126309 |
| 336 | MIR922 | 6.777316736 | 100126321 |
| 337 | MIR937 | 7.662517289 | 100126338 |
| 338 | MIR939 | 7.634854772 | 100126351 |
| 339 | MIR944 | 6.694329184 | 100126340 |
| 340 | MLLT11 | 5.154449055 | 10962 |
| 341 | MROH1 | 7.745504841 | 727957 |
| 342 | MROH5 | 7.745504841 | 389690 |
| 343 | MROH6 | 7.727063163 | 642475 |
| 344 | MRPL13 | 8.022130014 | 28998 |
| 345 | MRPL47 | 7.597971415 | 57129 |
| 346 | MTBP | 7.985246657 | 27085 |
| 347 | MTDH | 6.094974643 | 92140 |
| 348 | MTERFD1 | 5.689257722 | |
| 349 | MTSS1 | 8.796680498 | 9788 |
| 350 | MUC20 | 6.804979253 | 200958 |
| 351 | MUC4 | 6.804979253 | 4585 |
| 352 | MYC | 10.55786077 | 4609 |
| 353 | MYEOV | 5.726141079 | 26579 |
| 354 | MYNN | 8.114338405 | 55892 |
| 355 | NAALADL2 | 8.049792531 | 254827 |
| 356 | NACAP1 | 7.053941909 | |
| 357 | NAPRT1 | 7.727063163 | |
| 358 | NCALD | 6.934071 | 83988 |
| 359 | NCBP2 | 6.804979253 | 22916 |
| 360 | NCEH1 | 7.819271554 | 57552 |
| 361 | NDRG1 | 8.197325957 | 10397 |
| 362 | NDUFAF6 | 5.726141079 | 137682 |
| 363 | NDUFB5 | 7.597971415 | 4711 |
| 364 | NDUFB9 | 8.686030429 | 4715 |
| 365 | NIPAL2 | 6.159520516 | 79815 |
| 366 | NKD2 | 5.375749193 | 85409 |
| 367 | NLGN1 | 7.690179806 | 22871 |
| 368 | NMD3 | 5.30198248 | 51068 |
| 369 | NOTCH2 | 5.025357308 | 4853 |
| 370 | NOV | 7.920700784 | |
| 371 | NRBP2 | 7.717842324 | 340371 |
| 372 | NRROS | 6.814200092 | 375387 |
| 373 | NSMCE2 | 8.990318119 | 286053 |
| 374 | NUDCD1 | 6.823420931 | 84955 |
| 375 | OC90 | 8.390963578 | 729330 |
| 376 | ODF1 | 6.703550023 | 4956 |
| 377 | OPA1 | 6.685108345 | 4976 |
| 378 | OPLAH | 7.791609036 | 26873 |
| 379 | ORAOV1 | 6.22406639 | |
| 380 | OSR2 | 6.187183034 | 116039 |
| 381 | OSTN | 6.519133241 | 344901 |
| 382 | OTOL1 | 5.329644998 | 131149 |
| 383 | OXR1 | 6.915629322 | 55074 |
| 384 | PABPC1 | 6.897187644 | 26986 |
| 385 | PAK2 | 6.804979253 | 5062 |
| 386 | PARL | 7.496542185 | 55486 |
| 387 | PARP10 | 7.791609036 | 84875 |
| 388 | PCAT1 | 9.727985247 | 100750225 |
| 389 | PCAT2 | 9.718764408 | 103164619 |
| 390 | PCYT1A | 6.768095897 | 5130 |
| 391 | PDCD10 | 6.961733518 | 11235 |
| 392 | PDCD6 | 5.265099124 | 10016 |
| 393 | PDP1 | 5.320424159 | 54704 |
| 394 | PEX5L | 7.680958967 | 51555 |
| 395 | PHC3 | 7.966804979 | 80012 |
| 396 | PHF20L1 | 8.271092669 | 51105 |
| 397 | PI4KB | 5.108344859 | 5298 |
| 398 | PIGX | 6.814200092 | 54965 |
| 399 | PIGZ | 6.804979253 | 80235 |
| 400 | PIK3CA | 7.846934071 | 5290 |

| | Gene | AMP Freq % | entrez |
|---|---|---|---|
| 401 | PIP5K1A | 5.172890733 | 8394 |
| 402 | PKHD1L1 | 6.952512679 | 93035 |
| 403 | PLD1 | 7.717842324 | 5337 |
| 404 | PLEC | 7.763946519 | 5339 |
| 405 | PLEKHF2 | 5.633932688 | 79666 |
| 406 | PLEKHG4B | 5.265099124 | 153478 |
| 407 | POGZ | 5.126786538 | 23126 |
| 408 | POLR2H | 7.1461503 | 5437 |
| 409 | POLR2K | 6.73121254 | 5440 |
| 410 | POP1 | 6.131857999 | 10940 |
| 411 | POU5F1B | 10.22591056 | 5462 |
| 412 | PP13439 | 7.680958967 | |
| 413 | PPFIA1 | 5.827570309 | 8500 |
| 414 | PPM1L | 5.246657446 | 151742 |
| 415 | PPP1R16A | 7.634854772 | 84988 |
| 416 | PPP1R2 | 6.804979253 | 5504 |
| 417 | PRKCI | 7.893038266 | 5584 |
| 418 | PRPF3 | 5.006915629 | 9129 |
| 419 | PRUNE | 5.172890733 | |
| 420 | PSCA | 7.607192254 | 8000 |
| 421 | PSMB4 | 5.062240664 | 5692 |
| 422 | PSMD2 | 7.127708621 | 5708 |
| 423 | PSMD4 | 5.089903181 | 5710 |
| 424 | PTDSS1 | 5.71692024 | 9791 |
| 425 | PTK2 | 7.976025818 | 5747 |
| 426 | PTP4A3 | 7.745504841 | 11156 |
| 427 | PUF60 | 7.680958967 | 22827 |
| 428 | PVT1 | 10.49331489 | 5820 |
| 429 | PYCRL | 7.690179806 | |
| 430 | PYDC2 | 6.537574919 | 152138 |
| 431 | RAD21 | 7.699400645 | 5885 |
| 432 | RAD54B | 5.467957584 | 25788 |
| 433 | RBM12B | 5.228215768 | 389677 |
| 434 | RECQL4 | 7.634854772 | 9401 |
| 435 | RFC4 | 6.712770862 | 5984 |
| 436 | RFX5 | 5.089903181 | 5993 |
| 437 | RGS22 | 6.795758414 | 26166 |
| 438 | RHPN1 | 7.699400645 | 114822 |
| 439 | RIMS2 | 7.072383587 | 9699 |
| 440 | RN7SKP153 | 7.920700784 | 106481813 |
| 441 | RN7SKP155 | 8.547717842 | 106479164 |
| 442 | RN7SKP206 | 9.008759797 | 106479187 |
| 443 | RN7SKP222 | 6.602120793 | 106479193 |
| 444 | RN7SKP226 | 9.847856155 | 106480903 |
| 445 | RN7SKP234 | 7.395112955 | 106480368 |
| 446 | RN7SKP249 | 7.017058552 | 106480910 |
| 447 | RN7SKP265 | 7.570308898 | 106480914 |
| 448 | RN7SKP296 | 6.491470724 | 106481823 |
| 449 | RN7SKP298 | 6.666666667 | 106479228 |
| 450 | RN7SKP40 | 7.459658829 | 106479113 |
| 451 | RN7SKP52 | 7.561088059 | 106479118 |
| 452 | RN7SKP85 | 6.141078838 | 106479132 |
| 453 | RN7SL141P | 7.644075611 | 106480959 |
| 454 | RN7SL215P | 6.712770862 | 106480980 |
| 455 | RN7SL228P | 7.680958967 | 106480983 |
| 456 | RN7SL229P | 7.376671277 | 106479305 |
| 457 | RN7SL329P | 8.824343015 | 106479343 |
| 458 | RN7SL350P | 6.343937298 | 106481019 |
| 459 | RN7SL36P | 6.602120793 | 106479237 |
| 460 | RN7SL395P | 7.634854772 | 106481031 |
| 461 | RN7SL396P | 7.985246657 | 106479368 |
| 462 | RN7SL434P | 6.694329184 | 106480509 |
| 463 | RN7SL444P | 5.089903181 | 106479384 |
| 464 | RN7SL447P | 6.685108345 | 106479386 |
| 465 | RN7SL473P | 5.274319963 | 106479396 |
| 466 | RN7SL486P | 6.556016598 | |
| 467 | RN7SL563P | 7.035500231 | 106479427 |
| 468 | RN7SL590P | 8.934993084 | |
| 469 | RN7SL600P | 5.274319963 | 106481086 |
| 470 | RN7SL637P | 6.934071 | |
| 471 | RN7SL685P | 6.934071 | 106479473 |
| 472 | RN7SL703P | 7.588750576 | 106479479 |
| 473 | RN7SL738P | 6.694329184 | 106480695 |
| 474 | RN7SL773P | 6.804979253 | |
| 475 | RN7SL826P | 7.671738128 | 106479523 |
| 476 | RNA5SP147 | 7.487321346 | 100873414 |
| 477 | RNA5SP148 | 7.55186722 | 100873415 |
| 478 | RNA5SP149 | 7.505763024 | 106480437 |
| 479 | RNA5SP150 | 7.588750576 | 100873416 |
| 480 | RNA5SP151 | 7.524204703 | 100873417 |
| 481 | RNA5SP274 | 5.20055325 | 100873527 |
| 482 | RNA5SP275 | 6.721991701 | 100873528 |
| 483 | RNA5SP276 | 7.671738128 | 100873529 |
| 484 | RNA5SP277 | 8.003688336 | 100873530 |
| 485 | RNA5SP278 | 7.976025818 | 100873531 |
| 486 | RNF139 | 8.639926233 | 11236 |
| 487 | RNF168 | 6.721991701 | 165918 |
| 488 | RNF19A | 6.934071 | 25897 |
| 489 | RNU6ATAC24P | 6.804979253 | 106479554 |
| 490 | RNU6ATAC41P | 6.878745966 | 106481869 |
| 491 | RNU6ATAC8P | 6.620562471 | 106481161 |
| 492 | RNY4P25 | 5.089903181 | 100862665 |
| 493 | RNY4P5 | 8.289534348 | 100379299 |
| 494 | RNY5P3 | 7.662517289 | 100379664 |
| 495 | RPL22L1 | 7.644075611 | 200916 |
| 496 | RPL30 | 6.177962195 | 6156 |
| 497 | RPL35A | 6.611341632 | 6165 |
| 498 | RPL39L | 6.639004149 | 116832 |
| 499 | RPL8 | 7.634854772 | 6132 |
| 500 | RPRD2 | 5.172890733 | 23248 |

| | Gene | AMP Freq % | entrez |
|---|---|---|---|
| 501 | RRM2B | 6.758875058 | 50484 |
| 502 | RSPO2 | 6.749654219 | 340419 |
| 503 | RTP1 | 6.639004149 | 132112 |
| 504 | RTP2 | 6.602120793 | 344892 |
| 505 | RTP4 | 6.602120793 | 64108 |
| 506 | RUNX1T1 | 5.053019825 | 862 |
| 507 | SAMD12 | 8.049792531 | 401474 |
| 508 | SAMD7 | 8.022130014 | 344658 |
| 509 | SCARNA7 | 5.108344859 | 677767 |
| 510 | SCHIP1 | 5.080682342 | 29970 |
| 511 | SCNM1 | 5.09912402 | 79005 |
| 512 | SCRIB | 7.662517289 | 23513 |
| 513 | SCRT1 | 7.634854772 | 83482 |
| 514 | SCXA | 7.634854772 | |
| 515 | SCXB | 7.634854772 | |
| 516 | SDC2 | 5.781466113 | 6383 |
| 517 | SDHA | 5.265099124 | 6389 |
| 518 | SDHAP1 | 6.804979253 | 255812 |
| 519 | SEC62 | 8.012909175 | 7095 |
| 520 | SELENBP1 | 5.062240664 | 8991 |
| 521 | SEMA6C | 5.108344859 | 10500 |
| 522 | SENP2 | 7.026279391 | 59343 |
| 523 | SENP5 | 6.804979253 | 205564 |
| 524 | SERPINI1 | 6.980175196 | 5274 |
| 525 | SERPINI2 | 6.934071 | 5276 |
| 526 | SETDB1 | 5.283540802 | 9869 |
| 527 | SHANK2 | 5.707699401 | 22941 |
| 528 | SHARPIN | 7.736284002 | 81858 |
| 529 | SI | 6.177962195 | 6476 |
| 530 | SKIL | 7.819271554 | 6498 |
| 531 | SLA | 8.215767635 | 6503 |
| 532 | SLC12A7 | 5.375749193 | 10723 |
| 533 | SLC25A32 | 6.666666667 | 81034 |
| 534 | SLC2A2 | 7.625633933 | 6514 |
| 535 | SLC30A8 | 7.791609036 | 169026 |
| 536 | SLC39A4 | 7.634854772 | 55630 |
| 537 | SLC45A4 | 7.948363301 | 57210 |
| 538 | SLC51A | 6.758875058 | 200931 |
| 539 | SLC52A2 | 7.634854772 | 79581 |
| 540 | SLC6A18 | 5.375749193 | 348932 |
| 541 | SLC6A19 | 5.375749193 | 340024 |
| 542 | SLC7A14 | 7.810050715 | 57709 |
| 543 | SLC9A3 | 5.265099124 | 6550 |
| 544 | SLITRK3 | 6.196403873 | 22865 |
| 545 | SLURP1 | 7.597971415 | 57152 |
| 546 | SMC4 | 5.053019825 | 10051 |
| 547 | SMCO1 | 6.703550023 | 255798 |
| 548 | SNORA12\|E | 8.750576302 | 677800 |
| 549 | SNORA18\|E | 7.607192254 | 677805 |
| 550 | SNORA25\|E | 8.852005533 | 684959 |
| 551 | SNORA25\|E | 7.708621485 | 684959 |
| 552 | SNORA25\|E | 7.680958967 | 684959 |
| 553 | SNORA31\|E | 7.671738128 | 677814 |
| 554 | SNORA32\|E | 7.929921623 | 692063 |
| 555 | SNORA40\|E | 8.169663439 | 677822 |
| 556 | SNORA40\|E | 5.329644998 | 677822 |
| 557 | SNORA44 | 5.025357308 | 677825 |
| 558 | SNORA4\|EN | 7.496542185 | 619568 |
| 559 | SNORA4\|EN | 6.712770862 | 619568 |
| 560 | SNORA63\|E | 7.496542185 | 6043 |
| 561 | SNORA63\|E | 6.712770862 | 6043 |
| 562 | SNORA63\|E | 6.712770862 | 6043 |
| 563 | SNORA63\|E | 7.496542185 | 6043 |
| 564 | SNORA72\|E | 7.616413094 | 26775 |
| 565 | SNORA72\|E | 6.150299677 | 26775 |
| 566 | SNORA72\|E | 5.191332411 | 26775 |
| 567 | SNORA72\|E | 8.409405256 | 26775 |
| 568 | SNORA81\|E | 6.712770862 | 677847 |
| 569 | SNORA81\|E | 7.496542185 | 677847 |
| 570 | SNORD112 | 6.83264177 | 692215 |
| 571 | SNORD2\|EI | 6.712770862 | 619567 |
| 572 | SNORD5\|EI | 7.745504841 | 692072 |
| 573 | SNORD66\|I | 7.127708621 | 692107 |
| 574 | SNORD77\|I | 6.666666667 | 692197 |
| 575 | SNTB1 | 8.077455048 | 6641 |
| 576 | SNX27 | 5.016136468 | 81609 |
| 577 | SNX31 | 7.007837713 | 169166 |
| 578 | SOX2 | 7.570308898 | 6657 |
| 579 | SPAG1 | 6.860304288 | 6674 |
| 580 | SPATA16 | 7.644075611 | 83893 |
| 581 | SPATC1 | 7.791609036 | 375686 |
| 582 | SPTSSB | 5.338865837 | 165679 |
| 583 | SQLE | 8.805901337 | 6713 |
| 584 | SST | 6.611341632 | 6750 |
| 585 | ST3GAL1 | 8.197325957 | 6482 |
| 586 | ST6GAL1 | 6.703550023 | 6480 |
| 587 | STK3 | 6.380820655 | 6788 |
| 588 | SV2A | 5.006915629 | 9900 |
| 589 | SYBU | 6.851083449 | 55638 |
| 590 | TAF2 | 8.012909175 | 6873 |
| 591 | TARS2 | 5.145228216 | 80222 |
| 592 | TATDN1 | 8.667588751 | 83940 |
| 593 | TBC1D31 | 8.381742739 | 93594 |
| 594 | TBCCD1 | 6.795758414 | 55171 |
| 595 | TBL1XR1 | 7.561088059 | 79718 |
| 596 | TCTEX1D2 | 6.740433379 | |
| 597 | TERC | 8.095896727 | 7012 |
| 598 | TERT | 5.449515906 | 7015 |
| 599 | TFRC | 6.777316736 | 7037 |
| 600 | TG | 8.280313509 | 7038 |

| | Gene | AMP Freq % | entrez |
|---|---|---|---|
| 601 | THEM6 | 7.607192254 | 51337 |
| 602 | THPO | 7.1461503 | 7066 |
| 603 | TIGD5 | 7.708621485 | 84948 |
| 604 | TM4SF19 | 6.712770862 | 116211 |
| 605 | TMEM207 | 6.52835408 | 131920 |
| 606 | TMEM212 | 7.680958967 | 389177 |
| 607 | TMEM249 | 7.634854772 | 340393 |
| 608 | TMEM41A | 7.026279391 | 90407 |
| 609 | TMEM44 | 6.648224988 | 93109 |
| 610 | TMEM65 | 8.593822038 | 157378 |
| 611 | TMEM67 | 5.274319963 | 91147 |
| 612 | TMEM71 | 8.26187183 | 137835 |
| 613 | TMEM74 | 6.768095897 | 157753 |
| 614 | TMEM75 | 9.986168741 | |
| 615 | TMOD4 | 5.09912402 | 29765 |
| 616 | TNFAIP8L2 | 5.080682342 | 79626 |
| 617 | TNFRSF11B | 7.920700784 | 4982 |
| 618 | TNFSF10 | 7.883817427 | 8743 |
| 619 | TNIK | 7.791609036 | 23043 |
| 620 | TNK2 | 6.804979253 | 10188 |
| 621 | TONSL | 7.634854772 | 4796 |
| 622 | TOP1MT | 7.708621485 | 116447 |
| 623 | TP53INP1 | 5.587828492 | 94241 |
| 624 | TP63 | 6.869525127 | 8626 |
| 625 | TPPP | 5.265099124 | 11076 |
| 626 | TPRG1 | 6.924850161 | 285386 |
| 627 | TRA2B | 6.952512679 | 6434 |
| 628 | TRAPPC9 | 8.169663439 | 83696 |
| 629 | TRHR | 6.768095897 | 7201 |
| 630 | TRIB1 | 8.934993084 | 10221 |
| 631 | TRIM59 | 5.126786538 | 286827 |
| 632 | TRIP13 | 5.265099124 | 9319 |
| 633 | TRIQK | 5.182111572 | 286144 |
| 634 | TRMT12 | 8.630705394 | 55039 |
| 635 | TRPS1 | 7.800829876 | 7227 |
| 636 | TSNARE1 | 7.745504841 | 203062 |
| 637 | TSPYL5 | 5.781466113 | 85453 |
| 638 | TSTA3 | 7.699400645 | |
| 639 | TTC14 | 7.395112955 | 151613 |
| 640 | TUFT1 | 5.053019825 | 7286 |
| 641 | U3\|ENSG0( | 5.81834947 | |
| 642 | U3\|ENSG0( | 8.400184417 | |
| 643 | U4\|ENSG0( | 6.804979253 | |
| 644 | U8\|ENSG0( | 7.413554633 | |
| 645 | UBR5 | 6.73121254 | 51366 |
| 646 | UBXN7 | 6.740433379 | 26043 |
| 647 | UQCRB | 5.689257722 | 7381 |
| 648 | USP13 | 7.607192254 | 8975 |
| 649 | UTP23 | 7.717842324 | 84294 |
| 650 | UTS2B | 6.565237437 | 257313 |
| 651 | VPS13B | 6.786537575 | 157680 |
| 652 | VPS28 | 7.634854772 | 51160 |
| 653 | VPS72 | 5.09912402 | 6944 |
| 654 | VPS8 | 7.192254495 | 23355 |
| 655 | VWA5B2 | 7.136929461 | 90113 |
| 656 | WDR49 | 6.961733518 | 151790 |
| 657 | WDR53 | 6.694329184 | 348793 |
| 658 | WDYHV1 | 8.566159521 | |
| 659 | WISP1 | 8.169663439 | |
| 660 | XXYLT1 | 6.685108345 | 152002 |
| 661 | YEATS2 | 7.496542185 | 55689 |
| 662 | YWHAZ | 6.970954357 | 7534 |
| 663 | ZBBX | 6.915629322 | 79740 |
| 664 | ZC3H3 | 7.819271554 | 23144 |
| 665 | ZDHHC11 | 5.265099124 | 79844 |
| 666 | ZDHHC11B | 5.265099124 | 653082 |
| 667 | ZDHHC19 | 6.777316736 | 131540 |
| 668 | ZFAT | 8.197325957 | 57623 |
| 669 | ZFP41 | 7.699400645 | 286128 |
| 670 | ZFPM2 | 6.934071 | 23414 |
| 671 | ZHX1 | 8.520055325 | 11244 |
| 672 | ZHX2 | 8.36330106 | 22882 |
| 673 | ZMAT3 | 7.727063163 | 64393 |
| 674 | ZNF16 | 7.634854772 | 7564 |
| 675 | ZNF250 | 7.634854772 | 58500 |
| 676 | ZNF251 | 7.634854772 | 90987 |
| 677 | ZNF252P | 7.634854772 | 286101 |
| 678 | ZNF34 | 7.634854772 | 80778 |
| 679 | ZNF517 | 7.634854772 | 340385 |
| 680 | ZNF572 | 8.77823882 | 137209 |
| 681 | ZNF623 | 7.717842324 | 9831 |
| 682 | ZNF639 | 7.671738128 | 51193 |
| 683 | ZNF687 | 5.108344859 | 57592 |
| 684 | ZNF696 | 7.690179806 | 79943 |
| 685 | ZNF7 | 7.634854772 | 7553 |
| 686 | ZNF706 | 7.053941909 | 51123 |
| 687 | ZNF707 | 7.634854772 | 286075 |
| 688 | snoU13\|EN | 7.680958967 | |
| 689 | snoU13\|EN | 6.694329184 | |
| 690 | snoU13\|EN | 5.652374366 | |
| 691 | snoU13\|EN | 8.446288612 | |
| 692 | snoU13\|EN | 6.795758414 | |
| 693 | snoU13\|EN | 6.62978331 | |
| 694 | snoU13\|EN | 7.081604426 | |
| 695 | snoU13\|EN | 6.62978331 | |
| 696 | snoU13\|EN | 5.560165975 | |
| 697 | snoU13\|EN | 8.243430152 | |
| 698 | snoU13\|EN | 6.509912402 | |
| 699 | snoU13\|EN | 6.602120793 | |
| 700 | snoU13\|EN | 7.597971415 | |

| | Gene | AMP Freq % | entrez |
|---|---|---|---|
| 701 | snoU13\|EN | 7.579529737 | |
| 702 | snoU13\|EN | 6.998616874 | |
| 703 | snoU13\|EN | 7.533425542 | |
